# Multiplexed tumor profiling with generative AI accelerates histopathology workflows and improves clinical predictions

**DOI:** 10.1101/2023.11.29.568996

**Authors:** Pushpak Pati, Sofia Karkampouna, Francesco Bonollo, Eva Compérat, Martina Radic, Martin Spahn, Adriano Martinelli, Martin Wartenberg, Marianna Kruithof-de Julio, Maria Anna Rapsomaniki

## Abstract

Understanding the spatial heterogeneity of tumors and its links to disease is a cornerstone of cancer biology. Emerging spatial technologies offer unprecedented capabilities towards this goal, but several limitations hinder their clinical adoption. To date, histopathology workflows still heavily depend on hematoxylin & eosin (H&E) and serial immunohistochemistry (IHC) staining, a cumbersome and tissue-exhaustive process that yields unaligned tissue images. We propose the VirtualMultiplexer, a generative AI toolkit that translates real H&E images to matching IHC images for several markers based on contrastive learning. The VirtualMultiplexer learns from unpaired H&E and IHC images and introduces a novel multi-scale loss to ensure consistent and biologically reliable stainings. The virtually multiplexed images enabled training a Graph Transformer that simultaneously learns from the joint spatial distribution of several markers to predict clinically relevant endpoints. Our results indicate that the VirtualMultiplexer achieves rapid, robust and precise generation of virtually multiplexed imaging datasets of high staining quality that are indistinguishable from the real ones. We successfully employed transfer learning to generate realistic virtual stainings across tissue scales, patient cohorts, and cancer types with no need for model fine-tuning. Crucially, the generated images are not only realistic but also clinically relevant, as they greatly improved the prediction of different clinical endpoints across patient cohorts and cancer types, speeding up histopathology workflows and accelerating spatial biology.

## Introduction

Tissues are spatially organized ecosystems, where cells of diverse phenotypes, morphologies and molecular profiles coexist with non-cellular compounds and interact to maintain homeostasis [1]. Several tissue staining technologies are used to interrogate this intricate tissue architecture and identify morphological and molecular patterns linked to disease. Among these technologies, H&E staining is the undisputed workhorse, routinely used to assess aberrations in tissue morphology in histopathology workflows across diseases [2]. A notable example is cancer, where H&E staining is routinely used to reveal abnormal cell proliferation, nuclear shape, lymphovascular invasion and immune cell infiltration, among others. Complementary to the morphological information available *via* H&E staining, IHC [3], another staining technique routinely applied in histopathology laboratories, exploits antigen-antibody binding to detect and quantify the abundance of a single protein marker *in situ*. IHC visualizes the distribution and localization of specific markers within cell compartments (membranous, cytoplasmic and/or nuclear) and within their proper histological context, which is crucial for tumor subtyping, prognosis, and personalized treatment selection. As tissue re-staining in conventional IHC is limited, repeated serial tissue sections stained with different antibodies are required for in-depth tumor profiling. However, this is a time-consuming and tissue-exhaustive process, prohibitive in cases of limited tissue availability. At the same time, serial IHC staining yields unaligned, non-multiplexed tissue images occasionally of suboptimal quality in terms of tissue artifacts, and tissue unavailability may lead to missing stainings (Figure 1A). Recently, multiplexed imaging technologies (*e*.*g*., imaging mass cytometry - IMC [4], co-detection by indexing - CODEX [5], multiplexed ion beam imaging - MIBI [6]) have enabled the simultaneous quantification of dozens of markers on the same tissue, revolutionizing spatial biology [7]. Still, their high cost, cumbersome experimental process, tissue destructive nature, long turnaround time, and need for specialized personnel and equipment severely limit their clinical adoption.

**Fig. 1.**
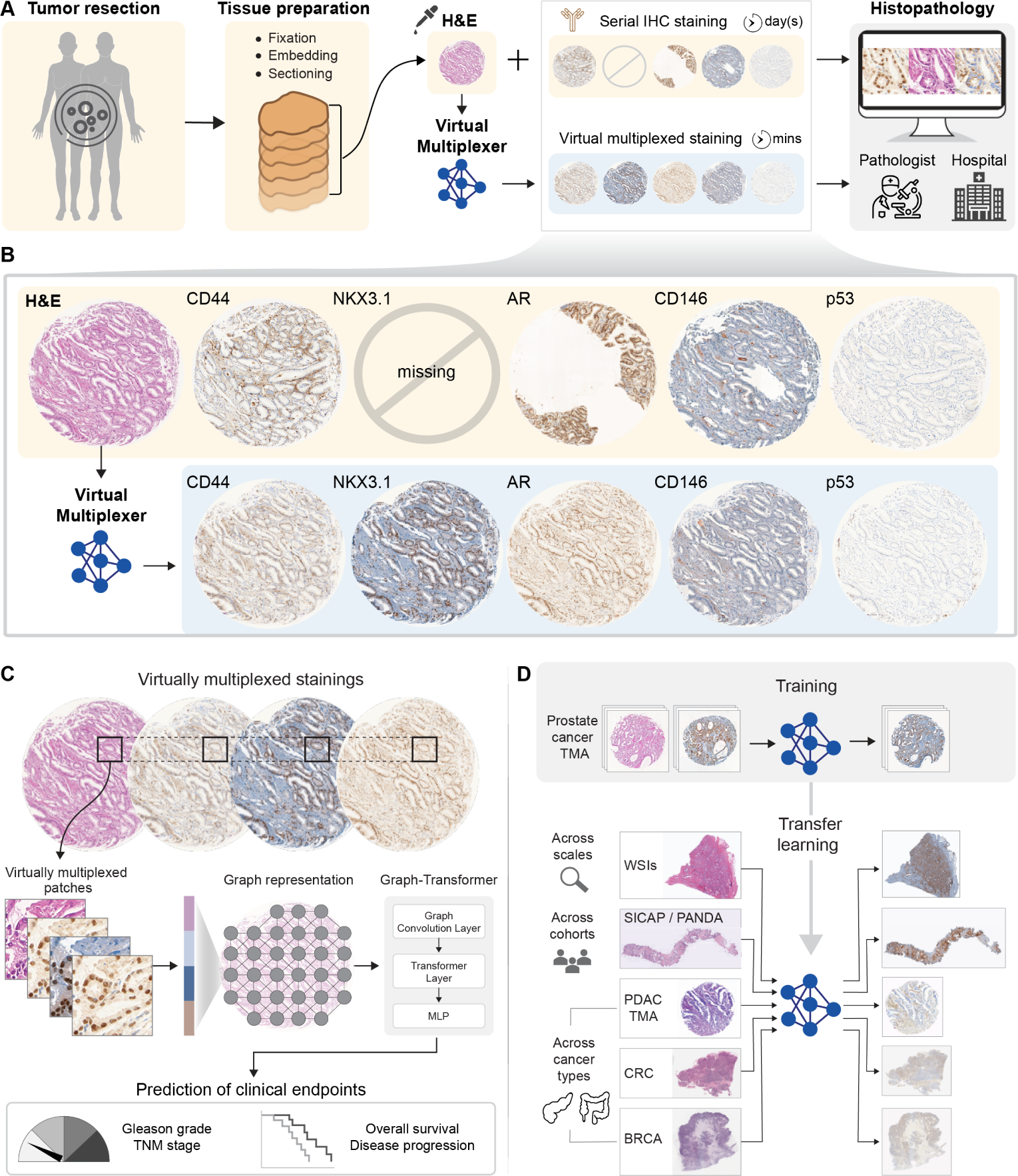
VirtualMultiplexer: a generative toolkit for synthesizing virtual multiplexed staining. **(A)** In a typical histopathology workflow, serial tissue sections from a tumor resection are stained with H&E and IHC to highlight tissue morphology and molecular expression of several markers of interest. This time-consuming and tissue-exhaustive process yields unpaired tissue slides that bear the technical risk of suboptimal quality in terms of missing stainings, tissue artifacts, and unaligned tissues. **(B)** To mitigate these issues, the VirtualMultiplexer uses generative AI to rapidly render, from a real input H&E image, consistent, reliable and pixel-wise aligned IHC stainings. **(C)** As the generated images are now virtually multiplexed, they are further exploited to train early-fusion Graph Transformers able to predict several clinically relevant endpoints. **(D)** The VirtualMultiplexer is transferable across image scales, patient cohorts and tissue types, accelerating clinical applications and discovery.

Virtual staining, *i*.*e*., the generation of artificially stained tissue images using generative AI models, has emerged as a promising cost-effective, easily accessible and rapid alternative that addresses above limitations [8, 9]. A virtual staining model exploits two sets of images - a *source* set and a *target* set - to learn the source-to-target appearance mapping [10, 11]. During inference, the trained model takes as input a source image and produces a virtually stained target image by simulating the target staining on the source. Initial virtual staining models were based on different flavors of generative adversarial networks (GANs) and operated under a *paired* setting, *i*.*e*., precisely aligned source and target images, which allowed them to directly optimize a pixel-wise loss between the virtual and real images [12]. Successful examples of paired models include translating label-free microscopy tissue images to H&E and specific stainings [13–16], H&E to special stains [17, 18], H&E to IHC [19, 20], and IHC to multiplex immunofluorescence [21]. However, as tissue re-staining is not routinely done in most cases, training paired models depends on aligning tissue slices via image registration, a time-consuming and error-prone process, which is often infeasible in practice because of substantial discrepancies even between consecutive slices. Additionally, as tissue architecture largely alters after the first set of slices, retrospective addition of new markers or multiplexing of several markers on a specific area/focal plane of interest is impossible. To circumvent these limitations, *unpaired* stain-to-stain (S2S) translation models have recently emerged, with early applications in translating from H&E to IHC [22–26] and special staining [27, 28] and from cryosections to Formalin-Fixed Paraffin-Embedded (FFPE) sections [29]. The vast majority of unpaired S2S translation models are inspired by CycleGAN [30]; they depend on an adversarial loss to preserve the source content (tissue architecture), and a cycle consistency loss to preserve the target style (staining pattern). Some employ additional constraints, *e*.*g*., domain-invariant content and domain-variant style [22], perceptual embeddings [24] or structural similarity [25].

However, an important limitation of CycleGAN-based models is that cycle consistency assumes a bijective mapping between the source and target domains [30], which does not necessarily hold for many S2S translation tasks. As a result, a persistent problem is staining unreliability, observed as incorrect mappings across domains, *e*.*g*., a positive signal from the source domain can get mapped to a negative signal from the target domain. To account for staining unreliability, recent works exploit expert annotations in the source and target domains to guide the translation. For example, Boyd *et al.* [26] translated H&E to Cytokeratin (CK) stained IHC using a region-based CycleGAN and expert annotations of positive and negative metastatic regions on the H&E images. Similarly, Liu *et al.* [25] translated H&E to Ki67 stained IHC by leveraging cancer or normal region annotations in both the H&E and IHC images. Although these approaches show promising results for these specific translation problems, acquiring such annotations is impractical when translating to several IHC markers, and infeasible even for experienced pathologists for more specialized translation tasks (*e*.*g*., identifying p53+ cells in H&E images). To circumvent the annotation challenge, Zheng *et al.* [31] recently introduced a semi-supervised approach, which however, again, depends on consecutive tissue sections and image registration. Consequently, there is a great need for S2S translation models that can learn from unpaired source and target images and preserve staining consistency without the need for consecutive tissue sections, image registration or extensive expert annotations on the source domain.

Regardless of the underlying modeling assumptions, an important limitation of the above S2S translation studies concerns model evaluation. As ground truth and virtually generated images are not pixel-wise aligned, S2S translation quality is typically quantified at a high-feature-level using inception-based scores [32]. However, these scores do not guarantee that the generated images accurately preserve complex and biologically meaningful spatial patterns [9]. To alleviate these concerns, a step in the right direction is expert qualitative assessment though pathological examination of the virtual images, employed by some studies [22, 24]. Still, as with all generative AI approaches, a persistent concern is the presence of hallucinations in virtually stained images that might otherwise appear realistic even to experienced pathologists. Ultimately, to ensure that virtually stained images do not only appear visually realistic but are also useful and relevant from a clinical standpoint, using them as input to downstream deep learning models to predict diagnostic or prognostic endpoints could provide an unbiased and convincing validation [9].

Here, we propose the VirtualMultiplexer, a generative toolkit that translates H&E images (source) to matching IHC images (target) for a variety of molecular markers, generating virtually multiplexed tissue images (Figure 1B). The VirtualMultiplexer is inspired by contrastive unpaired translation (CUT) [33], an appealing contrastive learning alternative to CycleGAN that achieves content preservation by maximizing the mutual information between target and source domains. Our toolkit does not depend on pixel-wise aligned H&E and IHC images during training and, in contrast to existing approaches, requires minimal expert annotations only on the target IHC domain. To ensure consistent and biologically relevant virtual stainings, VirtualMultiplexer introduces a novel architecture based on multi-scale constraints at the single-cell, cell-neighborhood and whole-image level that closely mimic human expert evaluation. Specifically, our multi-scale approach is designed to accurately capture the staining specificity at the individual cell level, while also ensuring content and style preservation at a cell neighborhood level and a global image level. We trained the VirtualMultiplexer on a prostate cancer tissue microarray (TMA) dataset containing unpaired H&E and IHC stainings for six clinically relevant nuclear, cytoplasmic, and membrane-targeted markers, and evaluated the generated images using quantitative image fidelity metrics, expert pathological assessment and visual Turing tests.

Importantly, to evaluate the clinical relevance of the generated images, we devised a Graph Transformer model that combined a Graph Neural Network and Vision Transformer (ViT) to simultaneously learn from the joint spatial distribution of several markers and predict clinically relevant endpoints (Figure 1C). As our toolkit is not limited by tissue availability or time constraints, we successfully transferred it across tissue image scales (TMAs to whole slide images (WSIs)), across two additional out-of-distribution large prostate cancer patient cohorts, and across tissue types (from prostate to pancreatic, breast and colorectal tissue) (Figure 1C). Our results suggest that the VirtualMultiplexer generates realistic multiplexed IHC stainings of high staining quality which are indistinguishable from real IHC stainings, outperforming existing methods. Importantly, using the generated datasets to train early fusion Graph Transformers surpassed in performance models trained with real unaligned data when predicting clinically relevant endpoints (*e*.*g*., survival status, tumor grade and stage) not only in the training cohort, but also in two independent prostate cancer patient cohorts and a pancreatic ductal adenocarcinoma (PDAC) cohort. Overall, the VirtualMultiplexer is a cost-effective, rapid, and easily accessible toolkit that can be readily used to generate virtual multiplexed imaging datasets of high quality, alleviate issues caused by missing modalities and tissue artifacts, improve the prediction of clinical endpoints and generalize across image scales, patient cohorts, and cancer types, with important implications in histopathology.

## Results

### VirtualMultiplexer: a generative toolkit for virtually multiplexed staining

The VirtualMultiplexer is a generative toolkit for unpaired S2S translation, trained on unpaired real H&E (source) and IHC (target) images. An overview of the model is presented in Figure 2 and a detailed description of its architecture and objective functions is provided in the Methods (Section 8). During training, each image is split into patches of size 256*×*256 pixels at 10*×* resolution, which are in turn fed into a generator network *G* that conditions on input H&E and IHC and learns to transfer the staining pattern, as captured from IHC images, to the tissue morphology, as captured by the H&E images. The generated IHC patches are then stitched together to create a final virtually stained IHC image (Figure 2A). During the S2S translation, we aim to ensure that the virtually stained IHC images are consistent with the real IHC images in terms of appearance, from the macroscopic to the microscopic scale. This implies that the generated IHC images should preserve the staining distribution of real IHC at a patch and tile scale, but also accurately learn the staining specificity of the different markers at the cellular scale. To achieve this, the model jointly optimizes three distinct loss functions, each one designed to preserve image consistency across different scales (Figure 2B). The neighborhood loss (1) ensures that the generated IHC patches are indistinguishable from real IHC patches and consists of an adversarial and a multilayer contrastive loss (Figure 2B), adopted from CUT [33]. The adversarial loss *L*_adv_ (1a) is a standard GAN loss [34], where real and virtual IHC patches are used as input to a convolutional neural network (CNN) patch discriminator network *D* which attempts to classify them as either real or virtual, eliminating style differences between real and virtual patches. The multilayer contrastive loss (1b) is based on a patch-level noise contrastive estimation loss [33] *L*_contrastive_ that aims to ensure that the content of corresponding real H&E and virtual IHC patches is preserved across multiple layers of *G*_enc_, *i*.*e*., the encoder of the generator *G*. The VirtualMultiplexer introduces two novel losses, namely a global and a local consistency loss (Figure 2B). The global consistency loss (2) uses a feature extractor *F*, a pre-trained VGG network [35], and enforces content consistency between the real H&E and the virtual IHC image (*L*_content_), and style consistency between the real and virtual IHC images (*L*_style_) at a tile level across multiple layers of *F*. In this way the model leverages a high-level sample information, i.e., the correspondence between real H&E and real IHC pairs, to ensure global tissue composition consistency and mitigate the macroscopic appearance variability. Finally, the local consistency loss (3) consists of a cell discriminator loss *L*_cellDisc_ and a cell classification loss (*L*_cellClass_) that together enable the model to capture a realistic appearance and staining pattern at the cellular level while alleviating the multi sub-domain mapping issue. This is achieved by leveraging prior knowledge on staining status via expert annotations, and training two separate networks: a cell discriminator *D*_cell_ that attempts to eliminate differences in the style of real and virtual cells, and a cell classifier *F*_cell_ that predicts the staining status and thus enforces staining consistency at a cell-level.

**Fig. 2.**
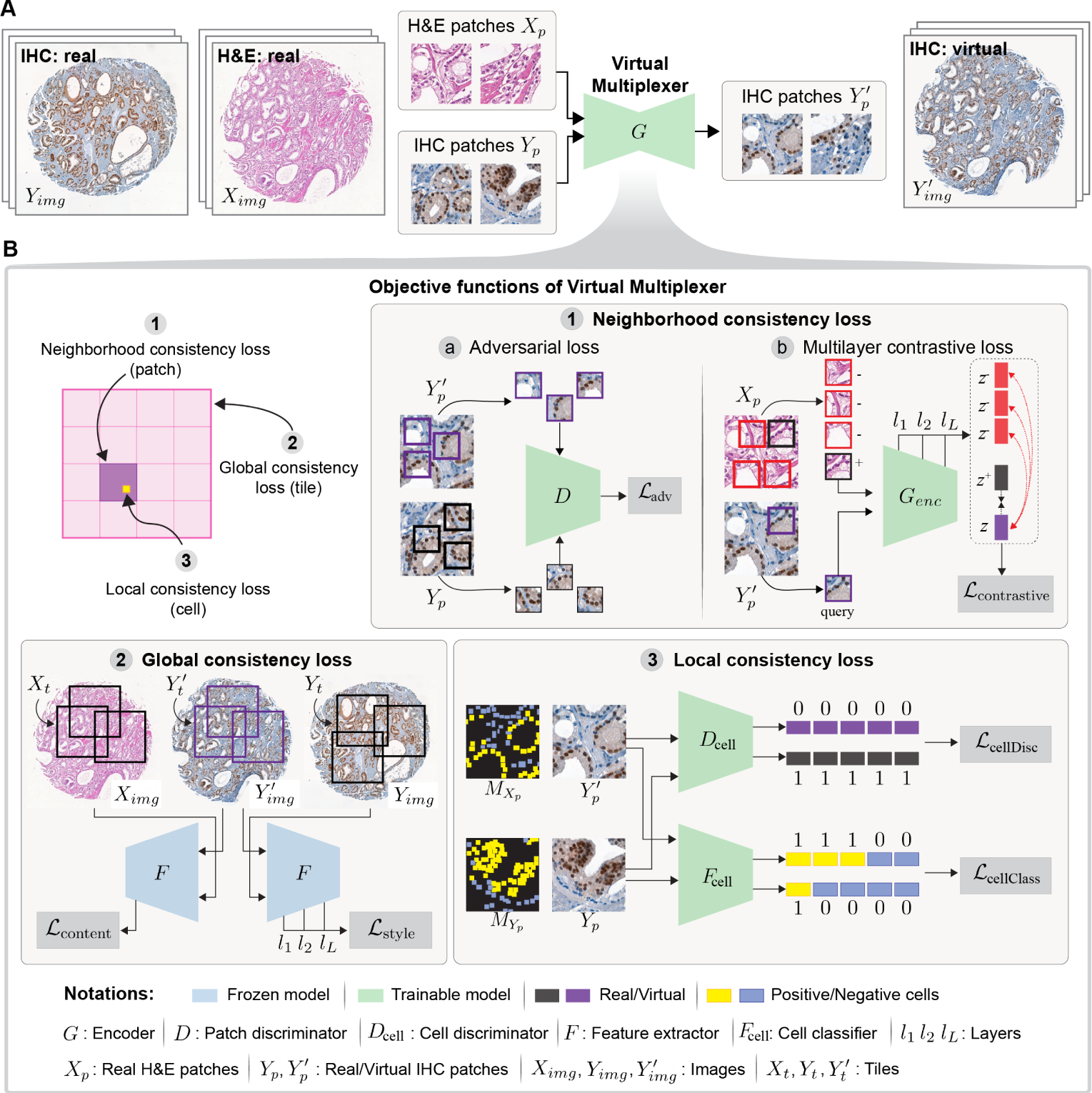
Overview of VirtualMultiplexer architecture. **(A)** The VirtualMultiplexer consists of a generator *G* that takes as input real unpaired H&E and IHC images and is trained to perform S2S translation by mapping the staining distribution of IHC onto H&E while preserving tissue morphology, ultimately generating virtually multiplexed synthetic IHC images only from input H&E images. **(B)** During training, the VirtualMultiplexer optimizes several losses that enforce consistent S2S translation at multiple scales, including a neighborhood consistency loss (1) that ensures indistinguishable translations at a neighborhood (patch) level, a global consistency loss (2) that ensures that the model accurately captures content and style constraints at a global tile-level, and a local consistency loss (3) that encodes biological priors on cell type classification and discriminator constraints at a cellular level.

### Performance assessment of the VirtualMultiplexer

We first assessed the performance of VirtualMultiplexer in terms of the quality of the generated IHC stainings. We trained the model on a cohort of prostate cancer TMAs from the European Multicenter Prostate Cancer Clinical and Translational Research Group (EMPaCT) [36–38] (Methods). The cohort contained unpaired H&E and IHC stainings from 210 patients with 4 cores per patient for six clinically relevant markers, namely androgen receptor (AR), NK3 Homeobox 1 (NKX3.1), CD44, CD146, p53 and ERG. The VirtualMultiplexer was used to generate a series of virtual IHC stainings (Figure 3C) that preserved the tissue morphology as seen in the real H&E image (Figure 3A) and the staining pattern as seen in the real IHC image (Figure 3B). Additional examples for two more TMA cores and all IHC markers are presented in the Appendix Figure A1. We quantitatively compared the results of the VirtualMultiplexer with four state-of-the-art unpaired S2S translation methods, namely CycleGAN [30], CUT [33], CUT with kernel instance normalization (KIN) [39], and AI-FFPE [29] using the Fŕechet Inception Distance (FID), an established metric used to assess the proximity of images created by a generative model to a set of target domain images [40]. The VirtualMultiplexer resulted in the lowest FID score across all markers (Figure 3D), with an average value of 29.2 (*±*3), which was consistently lower than the competing methods CycleGAN (49 *±* 6), CUT (35.8 *±* 4.5), CUT with KIN (37.8 *±* 2.3), and AI-FFPE (35.9 *±* 2.6). This result indicated that virtual stainings generated by the VirtualMultiplexer were the closest to the real IHC stainings in terms of distribution than any of the competing methods.

**Fig. 3.**
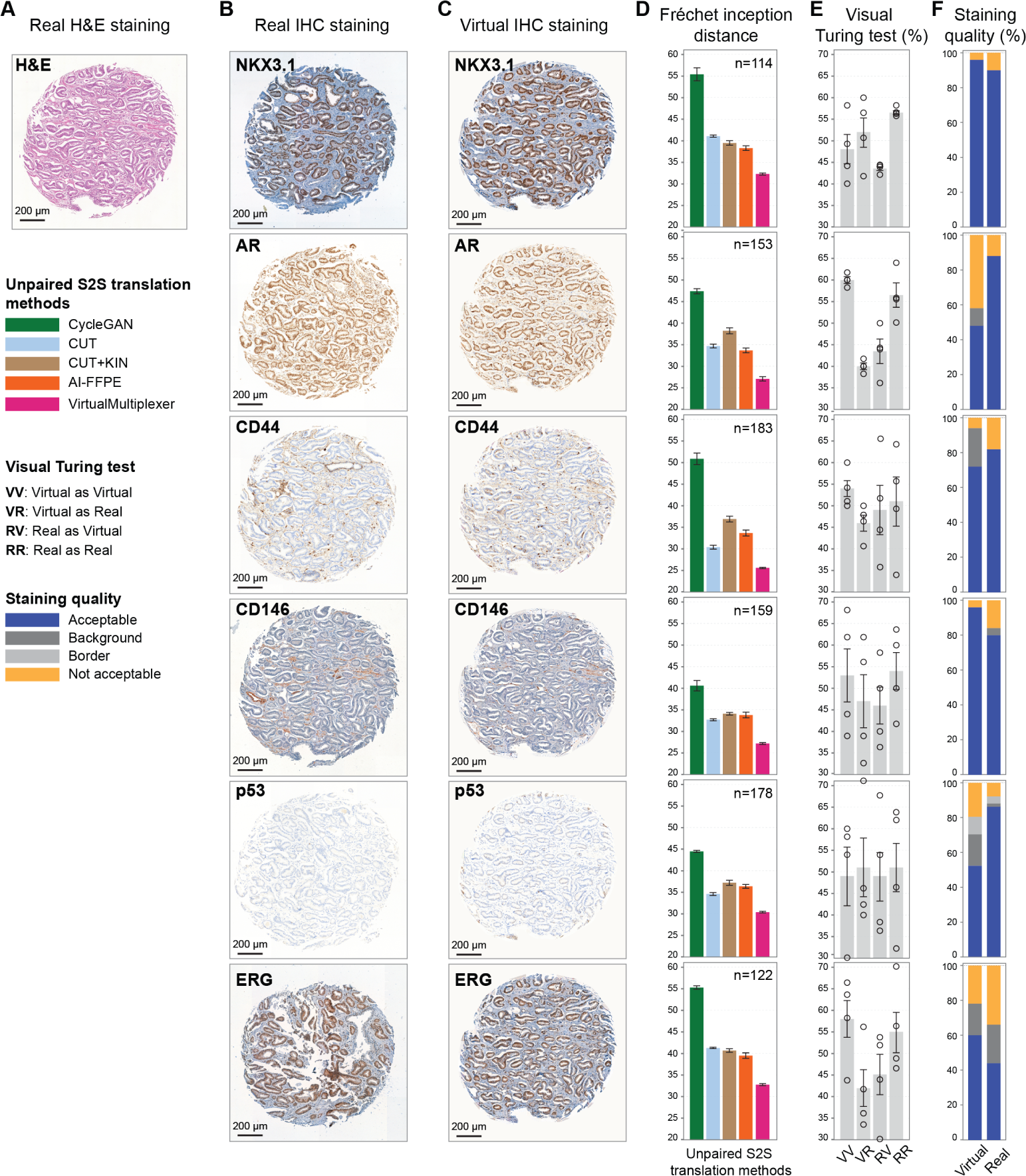
Performance evaluation of VirtualMultiplexer. **(A)** Example H&E core from the EMPaCT TMA. **(B)** Real, unpaired IHC-stained cores for different antibody markers corresponding to the H&E core in (A). **(C)** Virtually stained IHC cores, now paired with the H&E core in (A). **(D)** Comparison of the VirtualMultiplexer with state-of-the-art S2S models. Barplots and errorbars indicate the mean and standard deviation of the FID score from 3 independent runs of each model. Number of test samples used varies per marker and is reported in each subplot. **(E)** Results of the visual Turing test, where circles indicate the guess of each one of the four experts, and barplots and errorbars indicate the corresponding mean and standard variation. **(F)** Assessment of staining quality of the virtual and real stainings, performed on 50 real and 50 virtual images.

To further quantify the indistinguishability of real and synthetically stained images, we conducted a visual Turing test as follows: three independent evaluators with expertise in prostate tissue histopathology and one board-certified pathologist were shown 100 randomly selected patches per marker, with 50 of them originating from real and 50 from virtually stained IHC images, and were asked to classify each patch as virtual or real. We observed that our model was able to trick the experts, as it achieved an average sensitivity of 52.1% and specificity of 54.1% across all six markers (Figure 3E) (note that a random classification into real or virtual would correspond to a sensitivity and specificity of 50%). Last, we performed a staining quality assessment: we provided the pathologist 50 real and 50 virtual images from 50 TMA cores per IHC marker, revealing which are real and which are virtual, who in turn performed a qualitative assessment of the staining, as judged by overall expression levels, overall background, staining pattern, cell type specificity and subcellular localization (Figure 3F). Across all six markers, on average 70.7% of the virtually stained images reached an acceptable staining quality, as opposed to 78.3% of the real images. The results varied depending on the marker, with cores virtually stained for NKX3.1 and CD146 achieving the highest staining quality of 96%, surpassing even the real images. Conversely, virtually stained AR images had the lowest score of 46%, with an additional 10% exhibiting accurate staining but high background, and the remaining 42% rejected due to mostly heterogeneous staining, or falsely unstained cells. Background staining presented a challenge with CD44 and p53; the latter appeared to be further affected by border artifacts, *i*.*e*., presence of abnormally highly stained cells only in the border of the core, an artifact also occasionally present in real images. ERG achieved a higher staining quality assessment in virtual than in the real images, which both appeared to often face background issues. We concluded that, for most of the markers, the staining quality scores and the number of cores with staining artefacts (high background and/or border artefacts) were comparable in virtual versus real images.

Following these observations, we carefully examined the virtually generated images and assessed to which extend the VirtualMultiplexer captured the staining patterns of the real ones. The virtual images showed similar pattern and signal distribution to the real images for all six markers, with correct cell type and subcellular distribution (Figure 4A). For instance, AR+ and NKX3.1+ cells were evaluated as having correct distribution in the luminal epithelial compartment of the prostatic glands and nuclear localization. Furthermore, a few NKX3.1+ cells in stromal regions (possibly stroma-invading tumor cells) were correctly predicted in the virtually stained cores. Similarities in specific, matched areas between virtual and real IHC images were also assessed mainly for staining pattern and overall intensity levels. To this end, we specifically assessed that the virtual expression of markers indicative of tumor-specific molecular profile, such as loss of TP53 and ERG overexpression, did not largely deviate from the real IHC at the overall TMA core level (Figure 4A), which would be crucial for diagnostic applicability. Certain discrepancies in virtual versus real images were also found, such as non-specific signal in extra-cellular-matrix/stroma regions (NKX3.1, p53, ERG), false nuclear expression (CD44), and systematic lack of recognition of CD146+ vascular structures (Figure 4B). Nonetheless, the more pathologically relevant staining patterns were correctly reconstructed, as described in Figure 4A. We also performed an ablation study demonstrating the effects of training with different components of the objective function of the VirtualMultiplexer (Appendix Figure A2). Indeed, we observe that incorporating multi-scale loss terms, from the whole image to the cellular level, allows us to capture biologically consistent staining patterns that resemble the real IHC stainings. The mere imposition of the neighborhood consistency, the primary objective employed in competing methods, produces virtually stained tissue regions mimicking the overall staining distribution of real target IHC stainings, but clearly leads to staining unreliability, *e*.*g*., swapping of staining patterns between positive and negative cells. The introduction of the global consistency clearly mitigates this issue, and the further addition of the local consistency further optimizes the virtual staining at the cell-level.

**Fig. 4.**
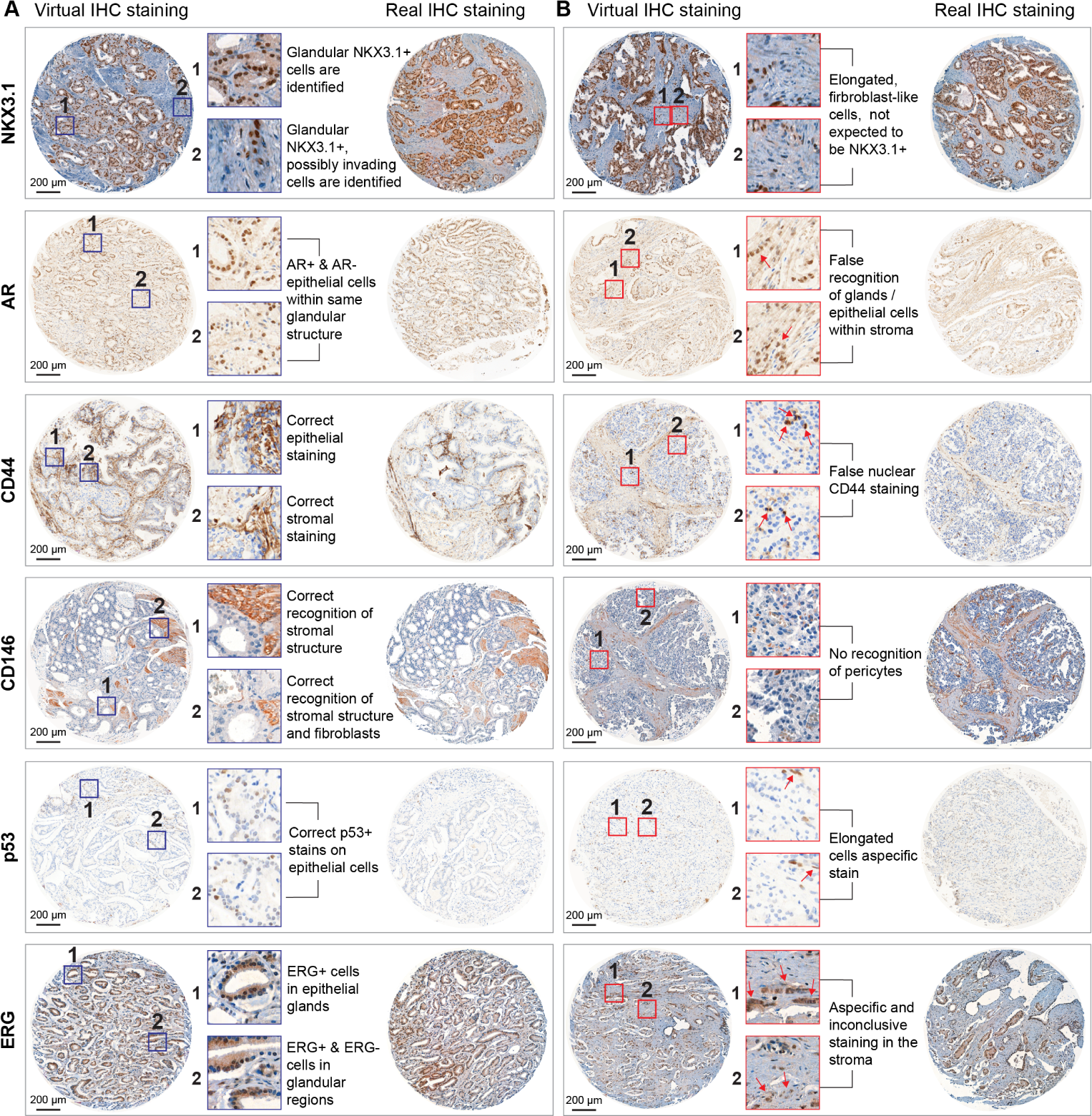
Visual quality assessment of virtually stained IHC images of the EMPaCT prostate cancer TMA. **(A)** Example virtual TMA cores across all 6 markers (left column) and selected zoomed in regions (middle column) that highlight accurate staining patterns. Real reference stainings for each markers are given on the right column. **(B)** Same as (A) but highlighting regions with inaccurate or inconclusive staining.

### Transfer learning from TMAs to WSIs

To assess how well the model can be transferred across imaging scales, we fed the TMA-trained VirtualMultiplexer with five out-of-distribution prostate tissue WSIs stained with H&E, and generated virtually stained IHC images for the NKX3.1, AR, and CD146 markers, as before. We then stained for the same markers by IHC on the direct serial sections, thus generating a real stained WSI that can be directly used to visually validate the model predictions. For marker NKX3.1 (Figure 5), we observed that the virtually stained images largely captured the staining appearance of the real ones, both in terms of specific glandular luminal cell identification (positive signal) (examples 1 and 2 in Figure 5 and Appendix Figure A3) and accurate non-annotation of stromal or vascular structures (absence of signal) (example 3 in Figure 5 and Appendix Figure A3). In minority, virtual images did not highlight the rarer NKX3.1+ cell population that are not part of the epithelial gland, but rather in the periglandular stroma (example 4 in Figure 5 and Appendix Figure A3). For both CD146 and AR, we observed staining intensity discrepancies between virtual and real images, which were more striking in the case of CD146 where the overall signal intensity and background is higher in the virtual versus the real images (Figure 5 and Appendix Figure A3). These discrepancies can be reasoned to the fact that the VirtualMultiplexer has been trained on TMA images that have a different staining distribution than the WSIs. This might lead to false interpretation of the marker expression levels at a first visual inspection. However, when evaluating at higher magnification, the staining pattern in the matching regions of real and virtual stainings was effectively correct, *e*.*g*., no glandular signal (example 5 in Figure 5) and appropriate stromal localization of CD146 (examples 6 and 7 in Figure 5) and nuclear localization of AR in luminal epithelial cells (example 1 in Appendix Figure A3). Lack of detection of vascular structures for CD146 was evident in both TMA cores and WSI (example 8 in Figure 5).

**Fig. 5.**
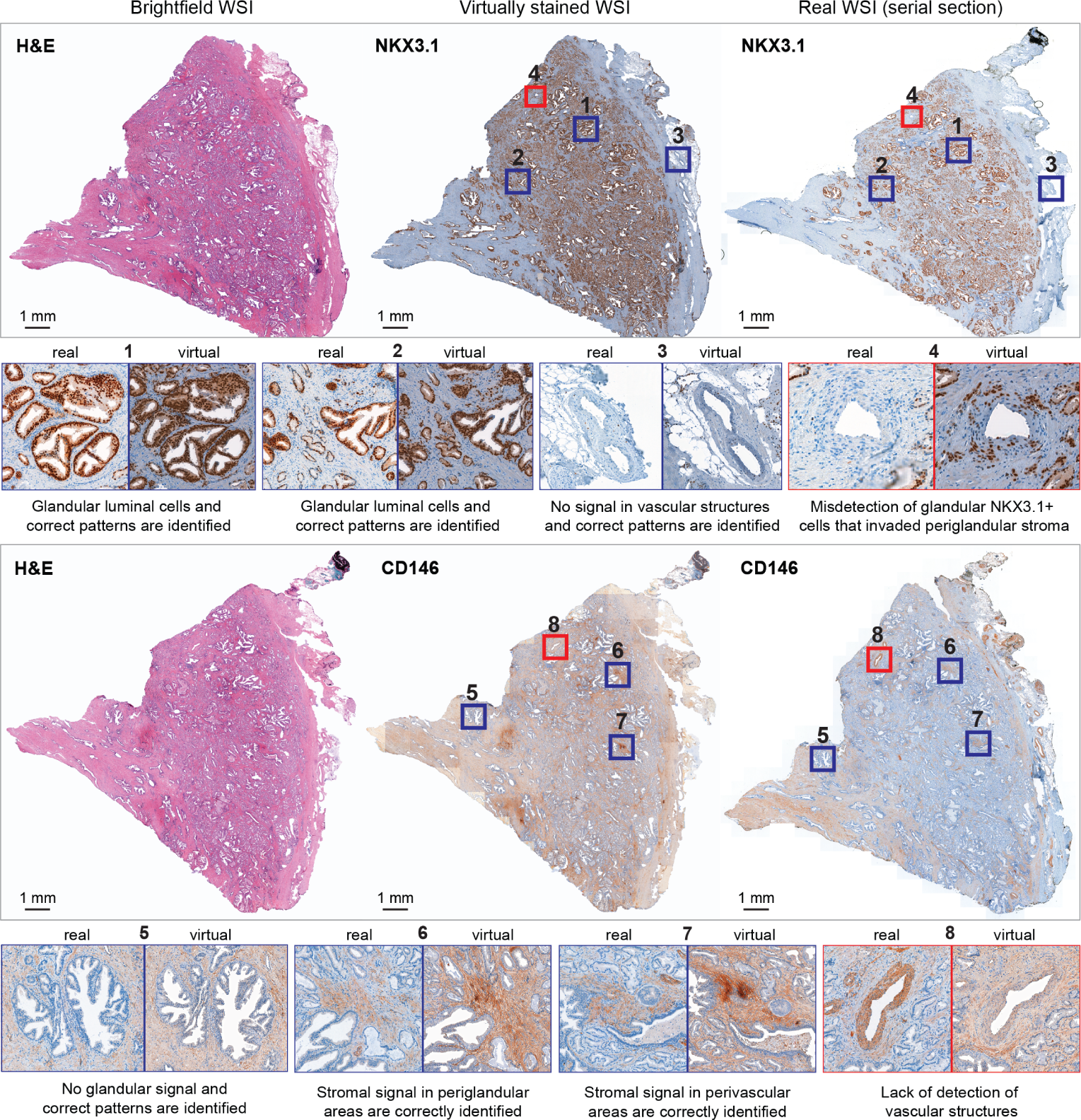
Transfer learning from TMA to WSIs of prostate cancer tissue. Example of H&E (left image), virtual IHC (middle image), and real IHC (right image) staining for NKX3.1 (top) and CD146 (bottom) of prostate cancer tissue WSIs. Blue-framed zoomed-in regions display accurate staining pattern. Red-framed zoomed-in regions display examples of virtual staining mis-predictions.

### The VirtualMultiplexer leads to improved clinical predictions

We further substantiated the utility of the generated virtual IHC stainings in augmenting the performance of AI models when predicting clinically relevant endpoints. To this end, we benchmarked the classification performance of AI models that were trained using real H&E, real IHC, or virtual IHC images. Specifically, we encoded the images as tissue-graph representations and employed a Graph-Transformer (GT) [41] to map the representations to downstream class labels. First, we extracted patches from the images and used a pretrained ResNet-50 network [42] to encode patch features (Figure 6A and Methods). The patches and their features formed the nodes and node representations of the tissue-graph, and the edges were formed using the spatial distribution of the patches (Figure 6B and Methods). The graph representation underwent graph convolutions to contextualize the node features of the local tissue neighborhood. Afterwards, the node features were pooled and fed to a transformer layer, trained to predict clinical endpoints. Depending on how the patch features were combined, we trained the GT model under the following three settings (Figure 6C): (i) a unimodal setting, where independent GT models were trained for each H&E and IHC marker, (ii) a multimodal late fusion setting, where the outputs of independent GT models were fused at the last embedding stage, and (iii) a multimodal early fusion setting, where the patch features were combined early in the tissue-graph and fed into the GT model. While the unimodal setting resulted in a separate prediction per marker, in both multimodal settings the patch features were combined, resulting in one prediction across all markers. However, in contrast to the late fusion multi-modal setting that necessitated the training of several GT models, in the early fusion case only one model that learned from the joint spatial distribution across all markers was trained, mimicking a multiplexed imaging scenario. With the exception of the early fusion setting that was only feasible for virtual images, we tested all three settings with both real and virtual images as input, resulting in a total of five different combinations (Figure 6D, legend).

**Fig. 6.**
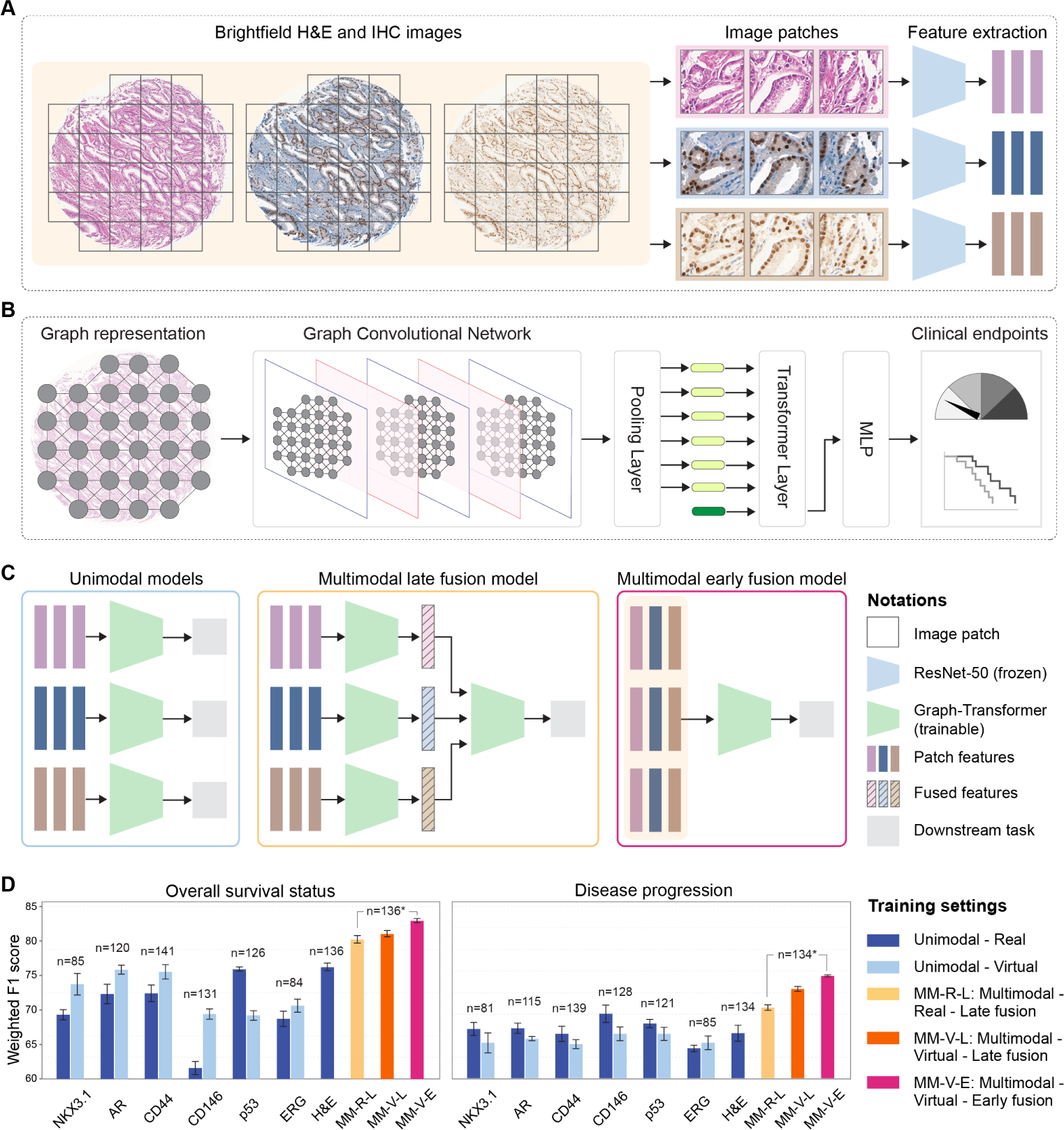
Prediction of clinically relevant downstream tasks with virtually multiplexed data. **(A)** Patch extraction and computation of patch features with a frozen ResNet-50 model (blue trapezoid). **(B)** Overview of the Graph-Transformer model, implemented by first constructing a patch-level graph representation, followed by a transformer that processes the graph representation to predict clinically relevant endpoints. **(C)** Training of Graph-Transformer models (green trapezoid) under three different settings, depending on the integration strategy. **(D)** Prediction results of overall survival status (left, 0: alive/censored, 1: prostate cancer related death) and disease progression (right, 0: no recurrence, 1: recurrence). Barplot colors indicate one of the five combinations of training setting and input data used (see legend). For each combination, barplot heights and errorbars indicate the mean and standard deviation of the weighted F1 score, as computed in the held-out test set from 3 independent runs with different initializations. The exact number of training samples used in each cases is given on the top of the barplots. * For all multimodal models, the reported number refers to the union across all markers.

We applied these settings to the EMPaCT dataset to predict two clinically relevant endpoints, namely the overall survival status and the disease progression of patients (Figure 6D). We note that in the EMPaCT dataset few IHC marker stainings are missing, leading to small discrepancies in the number of real IHC images available across all six markers. To ensure a fair comparison between real and virtual unimodal models, we matched the number of virtual IHC images to the number of available real IHC images, which implies that the dark and light blue barplots in Figure 6D are directly comparable. However, as H&E images were available for all patients, this issue did not affect the unimodal model trained on H&E images, which has a slight advantage over all other models in terms of number of samples used. Importantly, to compare all multimodal models, we again followed the same strategy of matching the number of virtual images used to the ones available in real data, and thus the last 3 bars in Figure 6D are also directly comparable. More details of the dataset distribution and the GT training are presented in Methods (Section 8). We observed that the unimodal models trained with virtual images are on par with unimodal ones trained with real images for both tasks, and the predictive performance of the unimodal models trained with virtual IHC images varied depending on the marker’s predictive ability towards the downstream task. In the case of overall survival status prediction, two interesting exceptions concern CD146 and p53: for CD146 the unimodal models trained on virtual data outperformed the ones trained on real data, which is in accordance to the previous observation that virtual CD146 images achieved a higher quality assessment than real CD146 images (Figure 3F). The opposite is true for p53: virtual p53 images were of lower quality than real p53 images, and the corresponding unimodal - virtual prediction models achieved a lower performance than the unimodal - real ones. However, these observations were not replicated for disease progression prediction, which appeared to be an overall harder prediction task. In both prediction tasks, the multimodal settings outperformed the unimodal H&E results, indicating the utility of combining information from complementary markers over individual stainings. Furthermore, the multimodal early fusion model trained with virtual images achieved the best weighted F1 score of 82.9% and 74.8% for overall survival status and disease progression, respectively, establishing the potential of multiplexed analysis via virtual staining for augmenting the efficacy of AI models. Overall, we concluded that using virtual images generated by the VirtualMultiplexer can boost the performance of state-of-the-art AI models for clinically relevant endpoints.

### Transferring the VirtualMultiplexer across patient cohorts and cancer types

We assessed the ability of the VirtualMultiplexer model to generalize to out-of-distribution data by employing two independent prostate cancer patient cohorts, namely SICAP [43] and PANDA [44], each one containing H&E stained needle biopsies with associated Gleason scores (details in Methods). We virtually stained the H&E images for four IHC markers, namely NKX3.1, CD146, AR and ERG, using the pre-trained VirtualMultiplexer on the EMPaCT dataset (example needle biopsy for SICAP in Figure 7A, additional examples for both SICAP and PANDA in Appendix Figure A4). The IHC markers were chosen due to their possible relevance towards Gleason score prediction. We observed that the virtual staining patterns of the IHC markers were overall correct and specific for each marker in terms of cell type and subcellular localization, with the only exception the occasional aspecific AR signal in the extracellular matrix areas. Other inconsistencies include the weak staining of interstitial tissue for CD146, and the heterogeneous staining of the glands for ERG. We also observed some recurring issues as in the quality assessment of the EMPaCT TMA (Figure 3), namely background (*e*.*g*., stromal background occasionally present in NKX3.1 and ERG), border and tiling artifacts (*e*.*g*., for CD146). Subsequently, we trained GT models under the previously described settings to predict the Gleason grade for both SICAP and PANDA datasets, shown in Figure 7B and C, respectively. We observe that the predictive performance of the unimodal models trained on virtual IHC stainings was close to or superior to the model using standalone H&E stainings for both the SICAP and PANDA datasets. Further improvement in Gleason score prediction was attained by the multimodal models built on the virtual IHC stainings. The early fusion model trained with the virtually multiplexed data achieved the best weighted F1 score of 61.4% and 72.3% for Gleason scoring on SICAP and PANDA, respectively, which significantly outperformed the H&E unimodal counterpart by 11.9% and 6.6% on SICAP and PANDA, respectively.

**Fig. 7.**
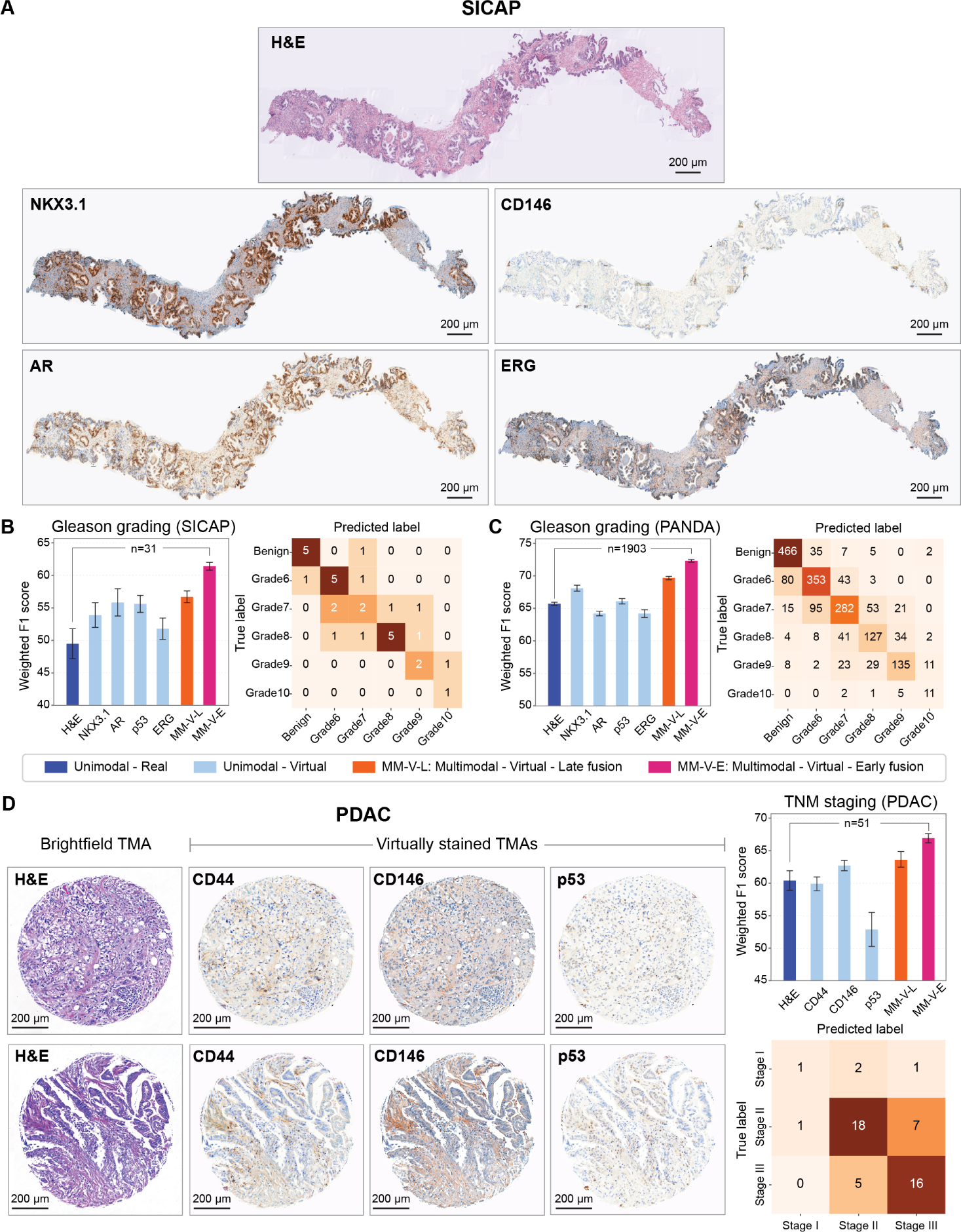
Transfer learning across scales, cohorts, and cancer types. **(A)** Real H&E needle biopsy of the SICAP dataset (top) and matching virtual IHC stainings across 4 IHC markers (bottom), as generated from the EMPaCT-trained VirtualMultiplexer. **(B)** Prediction results of Gleason grading for the SICAP test set in terms of weighted F1 score and confusion matrix. Barplots and errorbars as in Figure 3, confusion matrices correspond to the Multimodal - Virtual Early fusion model. Note that the setting unimodal - real (dark blue barplot) only includes training the model on H&E, as no real IHC data are available here. **(C)** Same as in (B) but for the PANDA dataset. **(D)** Virtual IHC staining of a PDAC TMA dataset with corresponding prediction of TNM staging.

Finally, we evaluated the generalization ability of the VirtualMultiplexer to images of other cancer types. To this end, we first applied the pre-trained VirtualMultiplexer on the EMPaCT dataset to a PDAC TMA that was unseen by the model and used the available H&E images to generate virtual IHC stainings for CD44, CD146 and p53 (Figure 7D), three markers with expected expression in pancreatic tissue. The generated images appeared overall realistic, with no means of discriminating whether they were virtually or actually stained. We observed that the CD44 and CD146 staining pattern in the virtual images was allocated, as expected, to the extracellular matrix of presented tissue spots, without major staining in the epithelial tissue part. For p53, we again observed overall proper staining allocation to the nuclei of epithelial cells with expected distribution, with no major staining of other compartments. To quantify the utility of the virtual stainings for downstream applications, we followed the same process as before to predict PDAC tumor, node and metastasis (TNM) stage, leading, again, to increased performance of models trained with virtually multiplexed data, concluding that virtually multiplexed data offers a performance advantage to prediction models.

We also applied the pre-trained VirtualMultiplexer to generate virtual IHC stainings for CD44 and CD146 from colorectal [45] and breast cancer [46] H&E-stained WSIs from The Cancer Genome Atlas (TCGA) [47]. Although the lack of normal tissue limited our ability to evaluate the staining quality in the generated images, we again observed an overall realistic virtual staining (Appendix Figure A5).

### The VirtualMultiplexer can greatly accelerate histopathology workflows

Lastly, we performed a runtime estimation of all components of the VirtualMultiplexer framework across imaging datasets of different scales, *i*.*e*., TMAs, needle biopsies and WSIs (Appendix Figure A6). We calculated that applying the trained VirtualMultiplexer on a single EMPaCT TMA core (6000*×*6000 pixels at 20X magnification-0.24*µ*m/pixel), an in-distribution sample, for one marker resulted in a total runtime of 2.81 seconds, and the same process for an out-of-distribution TMA core resulted in a runtime of 10.88 seconds, with the increase attributed to stain normalization. However, the stain normalization step is crucial as it alleviates the appearance disparity between the training EMPaCT H&E TMAs and the out-of-distribution samples, and allows for a faithful application of the VirtualMultiplexer to unseen datasets. The above result implies that virtual staining of a hypothetical TMA slide containing 250 out-of-distribution TMA cores for 6 markers would be feasible in *≈*65.8 minutes (preprocessing: *≈*9.9 seconds per core, virtual staining and post-processing: *≈*0.98 seconds per core and marker). Conversely, performing the IHC staining experiment for the same hypothetical TMA for 6 IHC markers could take an estimated time of approximately 1 day, when applied in a cutting-edge pathology laboratory using the latest protocols [48]. When applied in a biology lab that does not specialize in pathology, however, IHC staining could take up to 5 days per marker (sectioning: 1 day, staining: 2 days, slide drying: 1 day, imaging: 1 day), leading to a minimum of 5 days, if done simultaneously for all 6 markers, and more than 10 days, if performed mostly sequentially. Importantly, as our method scales linearly with the size of the tissue (TMA to WSI) and with the number of markers, similar time gains would be feasible for virtually staining needle biopsies and WSIs. We conclude that the VirtualMultiplexer leads to significant time gains when compared to a typical IHC staining, and can greatly accelerate histopathology workflows.

## Discussion

Virtual staining has emerged as a promising direction in histopathology, with early attempts to apply GAN-based approaches on unpaired datasets across different stain-to-stain translation tasks. However, staining inconsistency issues limit its applications in translating marker-specific stains such as IHC. In this work, we proposed the VirtualMultiplexer, a generative toolkit that can effectively translate H&E to IHC images for several markers, and is able to preserve staining consistency across image scales without requiring access to consecutive tissue sections, image registration or extensive expert annotations. This was achieved by proposing a novel architecture that includes the joint optimization of multi-scale loss functions that encode different biological priors to ensure biological consistency on a cellular, neighborhood, and global, whole-image scale. Our results indicated that the VirtualMultiplexer consistently out-performed in image fidelity state-of-the-art S2S translation methods and generated IHC images that were indistinguishable from the real ones to the expert human eye. Detailed histopathological evaluation suggested that the staining quality of the generated images was on par or even exceeded that of the real images in terms of staining pattern and distribution, with staining artifacts (*e*.*g*., high background, border effects) largely comparable in virtual versus real images for most markers. A thorough ablation study demonstrated that our novel local and global loss terms allowed us to mitigate staining unreliability and capture biologically consistent staining patterns, as opposed to solely using the adversarial and contrastive objectives employed in competing methods. When transferring the TMA-trained VirtualMultiplexer on prostate cancer WSIs and two unseen prostate cancer cohorts containing needle biopsies, we found that it generalized well to unseen prostate cancer images of different scales without any retraining or fine-tuning. Similar observations were made when generalizing on tissues of different origins, namely a PDAC TMA and TCGA breast and colorectal cancer WSIs.

While our results demonstrate a clear potential for the use of virtual multiplexed staining in histopathology, several limitations remain and can be addressed in future extensions of the model. Firstly, in some cases we observed elevated background levels in the EMPaCT TMA. Although this issue was also present in the real data, it appeared more pronounced for markers that had an overall more faint staining pattern (*e*.*g*., p53). More prominent background issues were present when we transferred the EMPaCT-trained model to the prostate cancer WSIs, which was expected considering that the EMPaCT IHC images and the prostate cancer WSIs were generated in two different institutions using different staining protocols and/or at different experiments and time points. Secondly, the patch-wise processing of the images induced tiling artifact in some cases, which is a well-known limitation of stain-to-stain translation approaches for large histopathology images [24, 39, 49]. In our algorithm, the tiling artifact was more pronounced at the border of a core, visible as patches of high staining intensity. One possible underlying cause of this effect is that, when the model receives as input an H&E patch at the border that contains very little tissue, the staining consistency losses “force” the model to stain the limited tissue with an overall higher intensity so as to match the staining distribution of the tissue-full patches that it has seen during training. Previous works [24, 39] have tried to address the tiling artifact, but it has been suggested that this caused less efficient translations [50]. As in our case the tiling artifact is observed in isolated, edge cases in the border, a straight-forward solution would be to simply discard a narrow border surrounding the tissue, as empirically done in regular IHC when border artifacts are present. Thirdly, certain discrepancies in staining specificity were occasionally observed, such as failing to stain CD146+ vascular structures and glandular NKX3.1+ cells that invaded periglandular stroma. This can be partially attributed to the fact that these patterns were observed more rarely in the training images, and can be mitigated by ensuring the inclusion of adequate representative examples in the labeling of the IHC images of the training set. Crucially, despite their limitations, the generated virtual images enabled the training of early fusion Graph-Transformer (GT) models, which consistently outperformed models trained on real data in the prediction on clinically relevant endpoints. This improvement was not only observed in the training TMA dataset across two prediction tasks, but also further confirmed on both independent prostate cancer cohorts, and the PDAC TMA cohort. In our experiments, we ensured that the multimodal early fusion GT models did not have an advantage in terms of number of samples used over the models trained with real data. At the same time, multimodal early fusion models had a much smaller parameter space in comparison to late fusion ones, as late fusion necessitated the training of as many GT models as the number of markers. This suggests that the higher model complexity of late fusion methods does not guarantee a performance improvement. A potential explanation of the observed performance improvement could be the quality of the generated images that are not affected by partially missing tumor areas or other artifacts occasionally found in real images. This explanation is corroborated by the fact that, for markers where virtual images were of higher quality than real images, the corresponding unimodal - virtual models out-performed the unimodal - real ones, and vice versa. A more likely explanation could be that, as early fusion models were able to learn from the joint spatial distribution of several markers on the same tissue at once, they were able to pick up multimodal spatial relationships at the cellular level, mimicking data generated by advanced multiplexed imaging technologies. This explanation is further supported by the fact that in the early fusion case, a single GT model proved to have more learning capacity than the integration of several equivalently potent ones.

In conclusion, the current work establishes the potential of virtual multiplexed staining across images of different scales, patient cohorts and tissue types, with important implications towards AI-assisted histopathology. For example, the VirtualMultiplexer could be directly used for data inpainting, *i*.*e*., filling out missing regions in a tissue image, or for sample imputation, *i*.*e*., generating from scratch missing samples in large patient cohorts. As IHC marker panels are not always standardized across labs, filling out the gaps via virtual multiplexed staining could open the door towards harmonizing datasets within or across research labs, which is particularly important in cases of archival tissue samples with limited availability [51, 52]. As a result, the VirtualMultiplexer could enable the generation of comprehensive patient cohorts that could be used for clinically relevant predictions. An equally important application of our work is the use of virtual multiplexed staining for pre-histopathological experimental design: generating a large collection of IHC stains *in silico* and training AI models could support marker selection for actual experimentation, significantly reducing costs and preserving precious tissue. To reach its full potential, future work will be crucial to establish the use of the VirtualMultiplexer in real-world settings. From a technical standpoint, the generated virtual multiplexed stainings can enable the development of foundational models for IHC, as they have been successfully developed for brightfield H&E images [53–55]. However, developing such models requires high volumes of data, which is potentially challenging to acquire for IHC. Virtual stainings can be beneficial to this end and can pave the way for multimodal foundational tissue characterization. Interestingly, the virtual multiplexed stainings can also be exploited as biologically conditioned data augmentations to boost the development and in turn the predictive performance of foundational models in histopathology applications. From an applications standpoint, although here we presented a first proof-of-concept for transferring the model across tissue types, more thorough evaluations are needed to solidify our encouraging results. Finally, although here we focused on H&E-to-IHC translation, as our method is stain-agnostic, straight-forward adaptations of our work for S2S translation across cutting-edge multiplexed imaging technologies (*e*.*g*., IMC, CODEX, MIBI) could significantly reduce their costs and find important applications in dataset harmonization or antibody panel selection and optimization. Our vision is that future extensions of our work could lead to the generation of an ever-growing and readily available dictionary of virtual stainers for IHC and beyond, that would be essentially limitless with respect to how many markers could be incorporated, surpassing in multiplexing ability even the most cutting-edge technologies and significantly accelerating spatial biology.

## Methods

### Datasets

The VirtualMultiplexer was trained using the European Multicenter High Risk Prostate Cancer Clinical and Translational research group (EMPaCT) TMA dataset; an independent subset of EMPaCT was used for internal testing. The VirtualMultiplexer was further evaluated in a zero-shot fashion, *i*.*e*., without any retraining or fine-tuning, on three external prostate cancer datasets, namely prostate cancer WSIs, SICAP [43] and PANDA [44] needle biopsies, on an independent PDAC dataset (PDAC TMAs) and on TCGA data from breast and colorectal cancer. In all cases, independent Graph-Transformers are trained and tested for individual datasets, by using both real and virtually stained samples, to address various downstream classification tasks. Details on all datasets used follow.

#### EMPaCT

The dataset contains TMAs from 210 primary prostate tissues, as part of the European Multicenter High Risk Prostate Cancer Clinical and Translational research group (EMPaCT) and the Institute of Tissue Pathology in Bern. The study followed the guidelines of the World Medical Association Declaration of Helsinki 1964, updated in October 2013, and was conducted after approval by the Ethics Committees of Bern (CEC ID2015-00128). For each patient, four cores were selected, with two of them representing a low Gleason pattern and the other two a high Gleason pattern. Consecutive slices from each core were stained with H&E and IHC using multiple antibodies against nuclear markers NK3 Homeobox 1 (NKX3.1) and Androgen receptor (AR), tumor markers Tumor Protein p53 (p53) and Erythroblast transformation-specific related gene (ERG) and membrane markers Cell Surface Glycoprotein (CD44) and Melanoma Cell Adhesion molecule (CD146/MCAM). TMA FFPE sections of 4 *µ*m were deparaffinized and used for heat-mediated antigen retrieval (citrate buffer, pH 6, Vector Labs or Tris-HCl pH 9). Sections were blocked for 10 min in 3% H_2_O_2_, followed by 30 minute room temperature incubation in 1% BSA in PBS–0.1%Tween 20. The following antibodies were used: anti-AR (Dako Agilent, M3562, 1:100 dilution), anti-NKX3.1 (Athena Enzyme Systems, 314, 1:200), anti-p53 (Dako Agilent, M7001, 1:800), anti-CD44 (Abcam, ab16728, 1:2000), anti-ERG (Abcam, ab133264, 1:500) and anti-CD146 (Abcam, ab75769 EPR3208, 1:500). Images were acquired using a 3D Histech Panoramic Flash II 250 scanner at 20*×* magnification (resolution 0.24*µ*m/pixel). The cores were annotated at patient-level by expert uro-pathologists with binary labels for overall survival status (0: alive/censored, 1: prostate cancer related death) and disease progression status (0: no recurrence, 1: recurrence). Clinical follow-up was recorded at a per-patient basis, with a maximum follow-up time of up to 12 years. For both the survival and disease progression clinical endpoints, the available data were imbalanced in terms of class distributions. Access information is possible upon request to the corresponding authors. The distribution of cores per clinical endpoint for the EMPaCT dataset is summarized in Table 1.

**Table 1.** Distribution of cores per clinical endpoint for the EMPaCT dataset.

| Overall survival status prediction |  |  |
| --- | --- | --- |
| #cores 0: Alive/censored | #cores 1: Prostate-cancer related death | #total cores |
| 472 | 205 | 677 |
| Disease progression prediction |  |  |
| #cores 0: No recurrence | #cores 1: Recurrence | #total cores |
| 568 | 109 | 677 |

#### Prostate cancer WSIs

Primary stage prostate cancer FFPE tissue sections (4 *µ*m) were deparaffinized and used for heat-mediated antigen retrieval (citrate buffer, pH 6, Vector Labs). Sections were blocked for 10 minutes in 3% H2O2, followed by 30 minute room temperature incubation in 1% BSA in PBS–0.1%Tween 20. The following primary antibodies were used: anti-CD146 (Abcam, ab75769 EPR3208, 1:500), anti-AR (Abcam, ab133273, EPR1535, 1:100) and anti-NKX3.1 (Cell Signaling, 83700T, 1:200). Secondary anti-rabbit antibody Envision HRP (DAKO, Agilent Technologies, Basel, Switzerland) for 30 minutes was used and signal detection was done using AEC substrate (DAKO, Agilent Technologies, Basel, Switzerland). Sections were counterstained with Hematoxylin and mounted with Aquatex). Images were acquired using a 3D Histech Panoramic Flash II 250 scanner at 20*×* magnification (resolution 0.24*µ*m/pixel).

#### SICAP

The dataset contains 155 H&E-stained WSIs from needle biopsies taken from 95 patients, split in 18,783 patches of size 512*×*512 [43]. The WSIs were reconstructed by stitching the patches. The WSIs were scanned at 40*×* magnification by Ventana iScan Coreo scanner and downsampled to 10*×* magnification. The WSIs were annotated by expert uro-pathologists for Gleason grades at the Hospital Cĺınico of Valencia, Spain.

#### PANDA

The dataset includes 5,759 H&E-stained needle biopsies from 1,243 patients at the Radboud University Medical Center, Netherlands [56], and 5,662 H&E stained needle biopsies from 1,222 patients at various hospitals in Stockholm, Sweden [57]. The slides from Radboud were scanned with a 3D Histech Panoramic Flash II 250 scanner at 20*×* magnification (resolution 0.24*µ*m/pixel) and were downsampled to 10*×*. The slides from Sweden were scanned with a Hamamatsu C9600-12 and an Aperio Scan Scope AT2 scanner at 10*×* magnification with a pixel resolution of 0.45202*µ*m and 0.5032*µ*m, respectively. The Gleason grades of the biopsies were annotated by expert uro-pathologists and were released as part of the Prostate cANcer graDe Assessment (PANDA) challenge [44]. We removed the noisy and inconspicuously labeled biopsies from the dataset, resulting in 4,564 and 4,988 biopsies from the Radboud and the Swedish cohorts, respectively (9,552 biopsies in total). The distribution of WSIs across Gleason grades for both SICAP and PANDA datasets are shown in Table 2.

**Table 2.** Distribution of WSIs across Gleason grades for both SICAP and PANDA datasets.

|  | Benign<br>0+0 | Grade 6<br>3+3 | Grade 7<br>3+4/4+3 | Grade 8<br>3+5/4+4/5+3 | Grade 9<br>4+5/5+4 | Grade 10<br>5+5 | Total |
| --- | --- | --- | --- | --- | --- | --- | --- |
| SICAP | 36 | 31 | 29 | 36 | 16 | 7 | 155 |
|  | Benign<br>0+0 | Grade 6<br>3+3 | Grade 7<br>3+4/4+3 | Grade 8<br>3+5/4+4/5+3 | Grade 9<br>4+5/5+4 | Grade 10<br>5+5 | Total |
| PANDA | 2603 | 2397 | 2327 | 1124 | 996 | 105 | 9552 |

#### PDAC

The PDAC TMA contained cancer tissue of 117 (50 female, 67 male) PDAC cases resected in a curative setting at the Department of Visceral Surgery of Inselspital Bern and diagnosed at Institute of Tissue Medicine and Pathology (ITMP) of the University of Bern between the years 2014 and 2020. The study followed the guidelines of the World Medical Association Declaration of Helsinki 1964, updated in October 2013, and was conducted after approval by the Ethics Committees of Bern (CEC ID2020-00498). All participants provided written general consent. The TMA contained three spots from each case (tumor front, tumor center, tumor stroma), leading to a total number of 351 tissue spots. Thirteen of these 117 cases were treated by neoadjuvant chemotherapy followed by surgical resection and adjuvant therapy, and the majority of the cases (104) were resected curatively and received adjuvant therapy. All cases were characterized comprehensively clinico-pathologically, including Tumour, Node, and Metastasis (TNM) stage during a master thesis of student Jessica Lisa Rohrbach at ITMP, supervised by Martin Wartenberg. All cases were UICC tumor stage I, stage II or stage III cases on pathologic examination, according to the Union for International Cancer Control (UICC) TNM Classification of Malignant Tumours, 8th edition [58]; the TMA did not comprise of UICC tumor stage IV cases. In all our analysis including the TNM prediction (Figure 7D), we excluded the thirteen neoadjuvant cases and considered only the 104 cases that received adjuvant therapy. The distribution of cores across the three TNM stages is reported in Table 3.

**Table 3.** Distribution of cores per TNM stage for the PDAC dataset.

| Stage I | Stage II | Stage III | Total |
| --- | --- | --- | --- |
| 24 | 121 | 115 | 260 |

#### TCGA

The dataset includes example H&E WSIs from breast (BRCA) and colorectal cancer (CRC) from The Cancer Genome Atlas (TCGA), available at the GDC data portal (https://portal.gdc.cancer.gov).

### Data Preprocessing

For all datasets used, we followed a tissue region detection and patch extraction preprocessing procedure. Specifically, the tissue region was segmented using the pre-processing tools in the HistoCartography library [59]. A binary tissue mask denoting the tissue and non-tissue regions was computed for each downsampled input image by iteratively applying Gaussian smoothing and Otsu thresholding until the mean of non-tissue pixels was below a threshold. The estimated contours of the denoted tissue and the cavities of tissue were then filtered depending on their area to generate the final segmentation mask. Subsequently, non-overlapping patches of size 256*×*256 were extracted from 10*×* magnification using the segmentation contours. The extracted H&E and IHC patches of the EMPaCT dataset were used for training and internal validation of the VirtualMultiplexer. The extracted H&E patches of all other unseen datasets (prostate cancer WSIs, SICAP, PANDA, PDAC, TCGA) were additionally first stain-normalized to mitigate the staining appearance variability with respect to the EMPaCT TMAs [60].

### VirtualMultiplexer Architecture

The VirtualMultiplexer is a generative AI toolkit that performs unpaired H&E-to-IHC translation. An overview of the model’s architecture is shown in Figure 2A. The VirtualMultiplexer is trained using two sets of images: source H&E images, denoted as *X*_img_ = *{x ∈ X}*, and target IHC images, denoted as *Y*_img_ = *{y ∈ Y}*. *X*_img_ and *Y*_img_ are unpaired images that originate from **different sections of the same TMA core** and thus belong to the same patient, but are pixel-wise unaligned, and thus unpaired. We train an independent one-to-one VirtualMultiplexer model for **each IHC marker** at a time. To train the VirtualMultiplexer, we use patches *X_p_* = *{x_p_ ∈ X*_img_*}* and *Y_p_* = *{y_p_ ∈ Y*_img_*}* extracted from a pair of images *X*_img_ and *Y*_img_, respectively. The backbone of the VirtualMultiplexer is a GAN-based generator *G*, specifically a Contrastive Unpaired Translation (CUT) [33] model, that consists of two sequential components, an encoder *G*_enc_ and a decoder *G*_dec_. Upon training, the generator takes as input a patch *x_p_* and generates a virtual patch 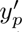, *i*.*e*., 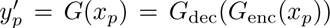. Finally, the virtually generated patches are stitched together to produce a final virtual image 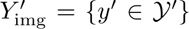. The VirtualMultiplexer is trained under the supervision of three levels of consistency objectives, namely local, neighborhood and global consistency (Figure 2B). The neighborhood consistency enforces effective staining translation at a patch-level, where a patch captures the neighborhood of a cell. We introduce additional global and local consistency objectives, operating at an image-level and cell-level, respectively, to further constrain the unpaired S2S translation and alleviate the stain-specific inconsistencies.

#### Neighborhood consistency

The neighborhood objective is a combination of an adversarial loss and a patch-wise multilayer contrastive loss, implemented as previously described in CUT [33] (Figure 2B, panel 1). Briefly, the adversarial loss dictates the model to learn to eliminate style differences between real and virtual patches, and the multilayer contrastive loss guarantees the content preservation at patch-level [61]. The adversarial loss is a standard GAN min-max loss [34], where the discriminator *D* takes as input real IHC patches *Y_p_* and IHC patches 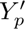 virtually generated by generator *G* and attempts to classify them as either real or virtual (Figure 2B, panel 1a). It is calculated as follows:

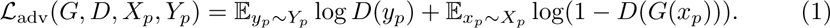

The patch-wise multilayer contrastive loss follows a noise contrastive estimation (NCE) concept as presented in [61, 62] and reused in [29, 33]. Specifically, it aims to maximize the resemblance between input H&E patch 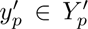 and corresponding virtually synthesized IHC patch 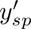 (Figure 2B, panel 1b). We first extract a *query* sub-patch 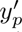 of size 64*×*64 from the target IHC domain patch 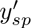 (purple square in panel 1b, Figure 2B) and match it to the corresponding sub-patch *x_sp_*, *i*.*e*., a sub-patch at the same spatial location as 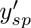 but from the H&E source domain patch *x_p_* (black square in panel 1b, Figure 2B). Since both sub-patches originate from the exact same tissue neighborhood, we expect that *x_sp_* and 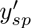 form a positive pair. We also sample *N* sub-patches 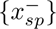 at different spatial locations from *x_p_* (red squares in panel 1b, Figure 2B), and expect that they form dissimilar, negative pairs with *x_sp_*. In a standard contrastive learning scheme, we would map *y_sp_*, *x_sp_*, and 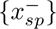 to a *d*-dimensional embedding space 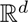 via *G_enc_* and project them to a unit sphere, resulting in *v, v*^+^ and 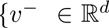, respectively, and then estimate the probability of a positive pair (*v, v*^+^) selected over negative pairs 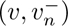*, ∀n ∈ N* as a cross-entropy loss with a temperature scaling parameter *τ*:

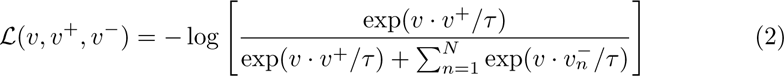

Here, we use a variation of the loss in Eq. (2), specifically a *patch-wise multilayer contrastive loss* that extends *L*(*v, v*^+^*, v^−^*) by computing it for feature maps extracted from *L*-layers of *G*_enc_ [29, 33]. This is achieved by passing the *L* feature maps of *x_p_* and 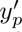 through a two-layer multilayer perceptron (MLP) *H_l_*, resulting in a stack of features 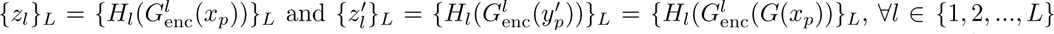, respectively. We also iterate over each spatial location *s ∈ {*1*, · · ·, S_l_}* and we leverage all *S*_l_*\s* patches as negatives, ultimately resulting in 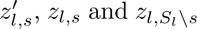 for the query, positive and negative sub-patches, respectively (purple, black and red boxes in Figure 2B, panel 1b). The final patch-wise multilayer contrastive loss is computed as:

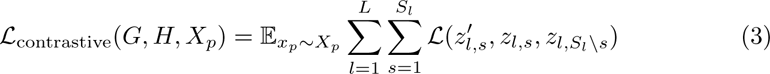

We also employ contrastive loss *L*_contrastive_(*G, H, Y_p_*) on patches *y_p_ ∈ Y_p_*, a domain-specific version of the identity loss [63, 64] that prevents the generator *G* from making unnecessary changes as proposed in [33]. Finally, the overall neighborhood consistency objective is computed as a weighted sum of the adversarial loss (1) and the multilayer contrastive loss (3) with regularization hyperparameter *λ*_NCE_:

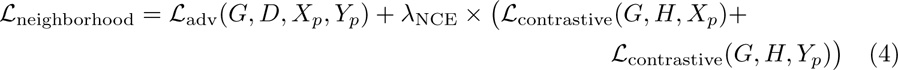

#### Global consistency

Inspired by seminal work in neural style transfer [65], this objective consists of two loss functions, a content loss *L*_content_ and a style loss *L*_style_ that together enforce biological consistency both in terms of tissue composition and staining pattern at the image (tile) level (Figure 2B, panel 2). Since the generated IHC images should be virtually paired to their corresponding input H&E image in terms of tissue composition, the content loss aims to penalize the loss in content between H&E and IHC images at a tile-level. First, real-patches *X_p_* and synthesized patches 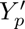 are stitched to create images *X*_img_ and 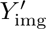, respectively, and corresponding tiles of size 1024 *×* 1024 are extracted (boxes in Figure 2B, panel 2), denoted as *X_t_* = *{x_t_ ∈ X*_img_*}* and 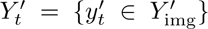, respectively. Then, the tiles are encoded by a pretrained feature extractor *F*, specifically VGG16 [35] pretrained on ImageNet [66]. The tile-level content loss at layer *l* of *F* is calculated as:

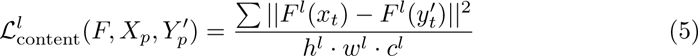

where, *h, w* and *c* are height, width, and channel dimensions of the feature map at *l*^th^ layer, respectively.

The style loss utilizes the synthesized image 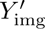 and the available real image *Y*_img_ to match the style or overall staining distribution between real and virtual IHC images. Since 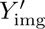 and *Y*_img_ do not have pixel-wise correspondence, large tiles 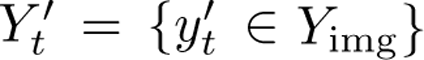 and *Y_t_* = *{y_t_ ∈ Y*_img_*}* are extracted at random such that each tile incorporates sufficient staining distribution. Next, 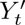 and *Y_t_* are processed by *F* to produce feature maps across multiple layers. The style loss is computed as:

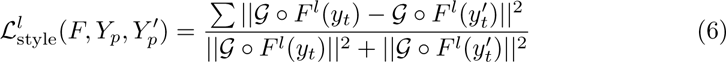

where *G* is the Gram matrix that measures the correlation between all the style in a feature map. The denominator is a normalization term which compensates for the under-or over-stylization of the tiles in a batch [67]. The overall global consistency loss is computed as:

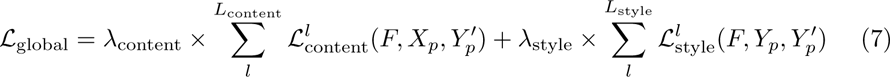

where *L*_content_ and *L*_style_ are the lists of the content and style layers of *F*, respectively, used to extract the feature matrices, and *λ*_content_ and *λ*_style_ are regularization hyperparameters for the respective loss terms.

#### Local consistency

The local consistency objective aims to enforce biological consistency at a local, cell-level and consists of two loss terms, namely a cell discriminator loss (*L*_cellDisc_) and a cell classification loss (*L*_cellClass_) (Figure 2B, panel 3). The cell discriminator loss is inspired by [26], and uses the cell discriminator *D*_cell_ to identify whether *a cell* is real or virtual, in the same way that the patch discriminator of Eq.(1) attempts to classify *patches* as real or virtual. *L*_cellDisc_ takes as input a real (*Y_p_*) and a virtual 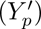 target patch and their corresponding cell masks (*M_Yp_* and 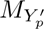, respectively), which include bounding box demarcation around the cells (Figure 2B, panel 3). *D*_cell_ comprises a feature extractor followed by a RoIAlign layer [68], and a final discriminator. The goal of *D*_cell_ is to output, *D*_cell_(*Y_p_, M_Yp_*) *→* **1** and 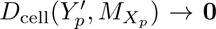, where **1** and **0** indicate real and virtual cells (indicated in black and purple, respectively, in Figure 2B, panel 3). The cell discriminator loss is defined as:

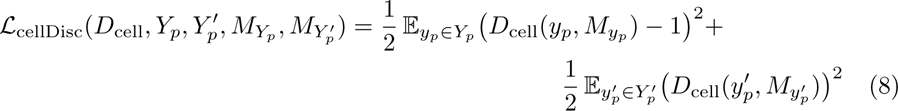

Although *D*_cell_ aims to enforce the generation of realistically looking cells, it is agnostic to their marker expression, as it does not explicitly capture which cells have a positive or a negative staining status. To account for this, we introduce an additional loss via a classifier *F*_cell_ that is trained to explicitly predict the cell staining status. This is achieved with the help of cell labels 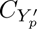 and *C_Yp_*, *i*.*e*., binary variables depicting the positive or negative staining status of a cell (indicated as 1: yellow and 0: blue boxes in Figure 2B, panel 3). The computation of cell masks and labels is described in detail in section **Cell masking and labeling of IHC images**. The cell-level classification loss can be easily computed as cross-entropy loss, calculated as:

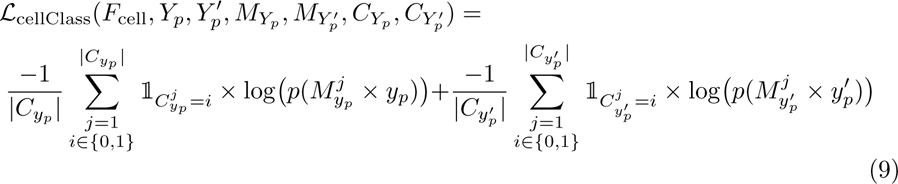

where, *|C_y_ |* and 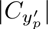 are the number of cells in *y_p_* and 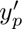, respectively, 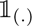 is the indicator function and *p*(.) is the cell-level probabilities predicted by *F*_cell_.

The overall local consistency loss is computed as:

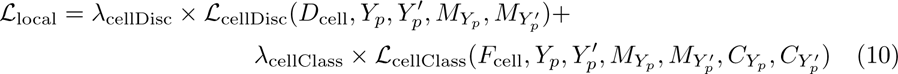

where *λ*_cellDisc_ and *λ*_cellClass_ are the regularization hyperparameteres for the cell discriminator and classification loss terms, respectively. Importantly, the local consistency loss can be easily generalized to any other cellular or tissue component (*e*.*g*., nuclei, glands) that might be relevant to other S2S translation problems, provided that corresponding masks and labels are available.

The complete objective function for optimizing VirtualMultiplexer is given as,

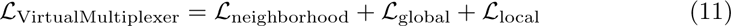

#### Cell masking and labeling of IHC images

As already discussed, the local consistency loss of Eq.11 needs as input cell masks *M_Xp_, M_Yp_* and cell labels *C_Xp_, C_Yp_*. However, acquiring these inputs manually for all patches across all antibodies is practically prohibitive, even for relatively small datasets. Automatic nuclei segmentation/detection using pretrained models (*e*.*g*., HoVerNet [69]) is a standard task for H&E images, but no such model exists for IHC images. To circumvent this challenge, we use an attractive property of the VirtualMultiplexer, *i*.*e*., its ability to synthesize virtual images that are pixel-wise aligned *in any direction* between the source and target domain. Specifically, we train a separate instance of the VirtualMultiplexer that performs IHC *→* H&E translation. The VirtualMultiplexer_IHC→H&E_ is trained using neighborhood consistency and global consistency objectives, as previously described. Once trained, it is used to synthesize a **virtual** H&E image 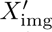 from a real IHC image *Y*_img_. At this point, we can leverage HoVerNet [69] to detect cell nuclei on real and virtual H&E images (*X*_img_ and 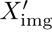) and simply transfer the corresponding cell masks (*M_X_*_img_ and 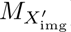) to their pixel-wise aligned IHC counterparts (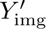 and *Y*_img_, respectively) to acquire 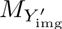 and *M_Y_*_img_. This “trick” eliminates the need to train individual cell detection models for each IHC antibody, and fully automates the cell masking process in the IHC domain. To acquire cell labels 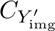 and *C_Y_*_img_, we use only region annotations in *Y*_img_, where the experts partially annotated areas as positive or negative stainings in a few representative images. Since IHC stainings are specialized in delineating positive or negative staining status, the annotation was easy and fast, and required approximately 2-3 minutes per image and per antibody marker. We also train cell detectors for the source and target domain, *i*.*e*., 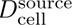, and 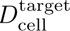, respectively. Provided the annotations, 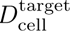 is trained as a CNN patch-classifier. The classifier predictions on *Y*_img_ combined with *M_Yp_* results in *C_Yp_*. The above region predictions on *Y*_img_ are transferred on to 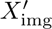. Afterwards, 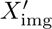 and the transferred annotatio 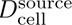 as a CNN patch-classifier. The classifier predictions on *X*_img_ combined with *M_Xp_* results in *C_Xp_*.

#### Implementation and training details

The architectural choices of the VirtualMultiplexer were set as follows: *G* is a ResNet [42] with 9 residual blocks; *D* is a PatchGAN discriminator [12]; *D*_cell_ includes four stride-2 feature convolutions followed by a RoIAlign layer and a discrimination layer; and *F*_cell_ includes four stride-2 feature convolutions and a 2-layer MLP. We use Xavier weight initialization [70], instance normalization [71], and a batch size of 1 image. We use Least Square GAN loss [72] for *L*_adv_. The model hyperparameters for the loss terms of the VirtualMultiplexer are set as: *λ*_NCE_ is 1 with temperature *τ* equal to 0.08, *λ*_content_ *∈ {*0.01, 0.1*}*, *λ*_style_ *∈ {*5, 10*}*, *λ*_cellDisc_ *∈ {*0.5, 1*}*, and *λ*_cellClass_ *∈ {*0.1, 0.5*}*. VirtualMultiplexer is optimized for 125 epochs using Adam optimizer [73] with momentum parameters *β*_1_ = 0.5 and *β*_2_ = 0.999. Different learning rates lr are employed for different consistency objectives, *i*.*e*., for neighborhood consistency lr*_G_* and lr*_D_* is set to 0.0002, for global consistency learning rate lr*_G_* is chosen from *{*0.0001, 0.0002*}*, and for local consistency learning rates lr*_D_*_cell_ and lr*_F_*_cell_ are chosen from *{*0.00001, 0.0001, 0.0002*}*. Among other hyperparameters, the number of tiles extracted per image to compute *L*_content_ and *L*_style_ is set to 8, the content layer in *F* is relu2 2, the style layers are relu1 2, relu2 2, relu3 3, relu4 3, and the number of cells per patch to compute *L*_cellDisc_ is set to 8.

### Graph-Transformer Architecture

The Graph-Transformer (GT) architecture, proposed by [41], fuses a Graph Neural Network and a vision transformer (ViT) to process histopathology images. The GNN first operates on a graph-structured representation of a histopathology image, where the nodes and edges of the graph denote patches and inter-patch spatial connectivity, and the nodes encode patch features extracted from a pre-trained deep learning model. Specifically, GT employs a graph convolution layer [74] to learn contextualized node embeddings through propagating and aggregating neighborhood node information. Subsequently, a ViT layer operates on the contextualized node features, leverages self-attention to weigh the significance of the nodes and aggregates the node information to render an image-level feature representation. Finally, an MLP maps the image-level features to a downstream image label. To note, histopathology images can have different spatial dimensions, therefore their graph-representations can have varying number of nodes. Also, the number of nodes can be very high when operating on giga-pixel sized WSIs. These two factors can potentially hinder the integration of the graph convolution layer to the ViT layer. To address these challenges, GT introduces a mincut pooling layer [75], which reduces the number of nodes into a fixed number of tokens while preserving the local neighborhood information of the nodes.

#### Implementation and training details

The architecture of GT follows the official implementation on github^1^. Each input image was cropped to create a bag of 256*×*256 non-overlapping patches at 10*×* magnification and background patches with non-tissue area ¿10% were discarded. The patches were encoded using ResNet50 [42] model pretrained on ImageNet dataset [66]. A graph representation was constructed using the patches with a 8-node connectivity pattern. The GT network consisted of one graph convolutional layer, and set the ViT layer configurations as, number of ViT blocks = 3, MLP size = 128, embedding dimension of each patch = 32, and number of multi-head attention = 8. The model hyperparameters were set as: number of clusters in mincut pooling = *{*50, 100*}*, Adam optimizer with initial learning rate of *{*0.0001, 0.00001*}*, a cosine annealing scheme for scheduling, and a mini-batch size of 8. The GT models were trained for 400 epochs with early stopping.

### Method Evaluation

#### Patch-level evaluation

We use the FID score [76] to compare the distribution of the virtual IHC patches with the distribution of the real IHC patches, as shown in Figure 3. The computation begins with projecting the virtual and the real IHC patches to an embedding space using InceptionV3 [76] model, pretrained on ImageNet [66]. The extracted embeddings are used to estimate multivariate normal distributions *N* (*µ_r_,* Σ*_r_*) for real data and *N* (*µ_s_,* Σ*_s_*) for virtual data. Finally, the FID score is computed as:

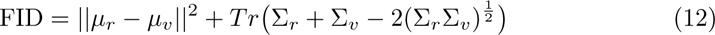

where, *µ_r_* and *µ_v_* are the feature-wise mean of the real and virtual patches, Σ*_r_* and Σ*_v_* are co-variance matrices for the real and virtual embeddings, and *Tr* is the trace function. A lower FID score indicates a lower disparity between the two distributions, thereby a higher staining efficacy of the VirtualMultiplexer. To ensure reproducibility, we ran each model three times with three independent initializations, and computed the mean and standard deviation for each model (barplot height and errorbar in Figure 3). We used a 70%-30% ratio to split the data in train and test sets, respectively. As for each marker a different number of IHC stainings were available in the EMPaCT data, the exact number of cores used per marker are given in Table 4:

**Table 4.** Distribution of cores per marker for training and testing the VirtualMultiplexer for the EMPaCT dataset.

|  | NKX3.1 | AR | CD44 | CD146 | p53 | ERG |
| --- | --- | --- | --- | --- | --- | --- |
| Train | 248 | 359 | 426 | 390 | 389 | 251 |
| Test | 114 | 153 | 183 | 159 | 178 | 122 |
| Total | 362 | 512 | 609 | 549 | 567 | 373 |

#### Image-level evaluation

We used a number of downstream classification tasks to assess the discriminative ability of the virtually stained IHC images on the EMPaCT, SICAP, PANDA and PDAC datasets. We further used these tasks to depict the utility of leveraging virtually multiplexed staining in comparison to standalone real H&E, real IHC, and virtual IHC staining. Specifically, provided the aforementioned images, we constructed graph representations as described in Section 8. Subsequently, Graph-Transformers [41] were trained under uni-modal and multi-modal settings using both real and virtually stained images, and evaluated on held-out independent test dataset. The final classification scores were reported using a weighted F1-metric, where a higher score depicts a better classification performance, thereby a higher discriminative power of the utilized images. As before, we ran each model three times with three independent initializations, and computed the mean and standard deviation for each model (barplot heights and errorbars in Figures 6 and 7). In all cases, we used a 60%-20%-20% ratio to split the data in train, validation and test sets, respectively. The exact number of train, validation and test samples used per task, marker and training setting in the EMPaCT dataset are given in Table 5.

**Table 5.** Distribution of cores for training, validation, and testing of Graph-Transformers per downstream task on the EMPaCT dataset, for all unimodal (first 7 columns) and multimodal models. * The number of cores reported under multimodal setting refers to the union across all markers used.

| Task: Overall survival status prediction |  |  |  |  |  |  |  |  |
| --- | --- | --- | --- | --- | --- | --- | --- | --- |
|  | NKX3.1 | AR | CD44 | CD146 | p53 | ERG | H&E | Multimodal* |
| Train | 232 | 333 | 397 | 343 | 377 | 230 | 407 | 407 |
| Validation | 74 | 110 | 133 | 123 | 120 | 82 | 134 | 134 |
| Test | 85 | 120 | 141 | 131 | 126 | 84 | 136 | 136 |

| Task: Disease progression prediction |  |  |  |  |  |  |  |  |
| --- | --- | --- | --- | --- | --- | --- | --- | --- |
|  | NKX3.1 | AR | CD44 | CD146 | p53 | ERG | H&E | Multimodal* |
| Train | 223 | 341 | 405 | 348 | 384 | 225 | 408 | 408 |
| Validation | 87 | 107 | 127 | 121 | 121 | 86 | 135 | 135 |
| Test | 81 | 115 | 139 | 128 | 128 | 85 | 134 | 134 |

For the SICAP, PANDA and PDAC datasets, the exact number of samples used in the train, validation and test splits coincide for all unimodal and multimodal models of Figure 7 and are reported in Table 6.

**Table 6.**
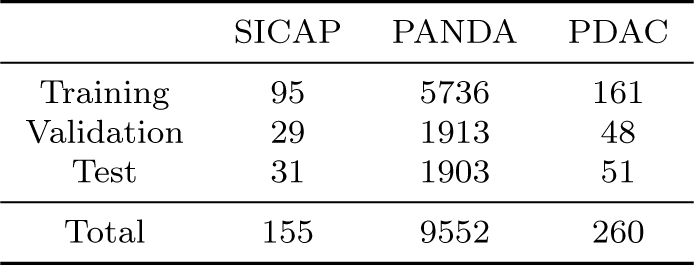
Number of available training, validation and test samples for the SICAP, PANDA and PDAC datasets.

|  | SICAP | PANDA | PDAC |
| --- | --- | --- | --- |
| Training | 95 | 5736 | 161 |
| Validation | 29 | 1913 | 48 |
| Test | 31 | 1903 | 51 |
| Total | 155 | 9552 | 260 |

### Computational Hardware and Software

The image datasets were preprocessed on POWER9 CPUs (Central Processing Units) and one NVIDIA Tesla A100 GPU (Graphics Processing Unit) using the Histocartography library [59]. The deep learning models were trained on NVIDIA Tesla P100 GPUs using PyTorch (version 1.13.1) [77] and PyTorch Geometric (version 2.3.0) [78]. The entire pipeline was implemented in Python (version 3.9.1).

## Data Availability

The main dataset used to support this study (EMPaCT) has been deposited in Zenodo, together with the prostate cancer WSIs (doi: 10.5281/zenodo.10066853). The SICAP dataset is available at Mendeley data (doi: 10.17632/9xxm58dvs3.1). The PANDA dataset is available at the Kaggle website (https://www.kaggle.com/ c/prostate-cancer-grade-assessment/data). The TCGA WSIs (BRCA and CRC) are available at the GDC data portal (https://portal.gdc.cancer.gov). The PDAC dataset is available upon reasonable request from the authors. All clinical data associated with the EMPaCT and PDAC patient cohorts cannot be shared owing to patient-confidentiality obligations.

## Author contributions

P.P. conceived and implemented the model. P.P., A.M. and M.A.R. designed and performed computational analyses. S.K., F.B. and M.R. performed the experiments. S.K., F.B., E.C., M.W. and M.K.d.J. assessed the virtual stainings. P.P., S.K., F.B., A.M. and M.A.R compiled the figures. M.S., M.W. and M.K.d.J. contributed materials for the experiments. P.P., S.K., F.B. and M.A.R. wrote the paper with inputs from all authors. M.K.d.J. and M.A.R. were responsible for the overall planning and supervision of the project.

## Code Availability

All source code of the VirtualMultiplexer is available under an open-source license in https://github.com/AI4SCR/VirtualMultiplexer.

## Acknowledgments

We would like to thank Guillaume Jaume, Jannis Born and Mara Graziani for constructive comments, discussions and suggestions. The results published here are in part based upon data generated by the TCGA Research Network: https://www. cancer.gov/tcga. This work was supported by the Swiss National Science Foundation (SNSF) Sinergia Grant (CRSII5 202297) to M.A.R. and M.K.dJ. The PDAC TMA construction has taken place at the Translational Research Unit (TRU) Platform of ITMP (https://www.ngtma.com/) in the setting of a grant by the Foundation for Clinical-Experimental Cancer Research Bern to M.W.

## Footnotes

1 A Graph-Transformer for Whole Slide Image Classification: https://github.com/vkola-lab/tmi2022

