## Supplementary Figures for "Multiplexed tumor profiling with generative AI accelerates histopathology workflows and improves clinical predictions"

### Appendix A Supplementary Figures

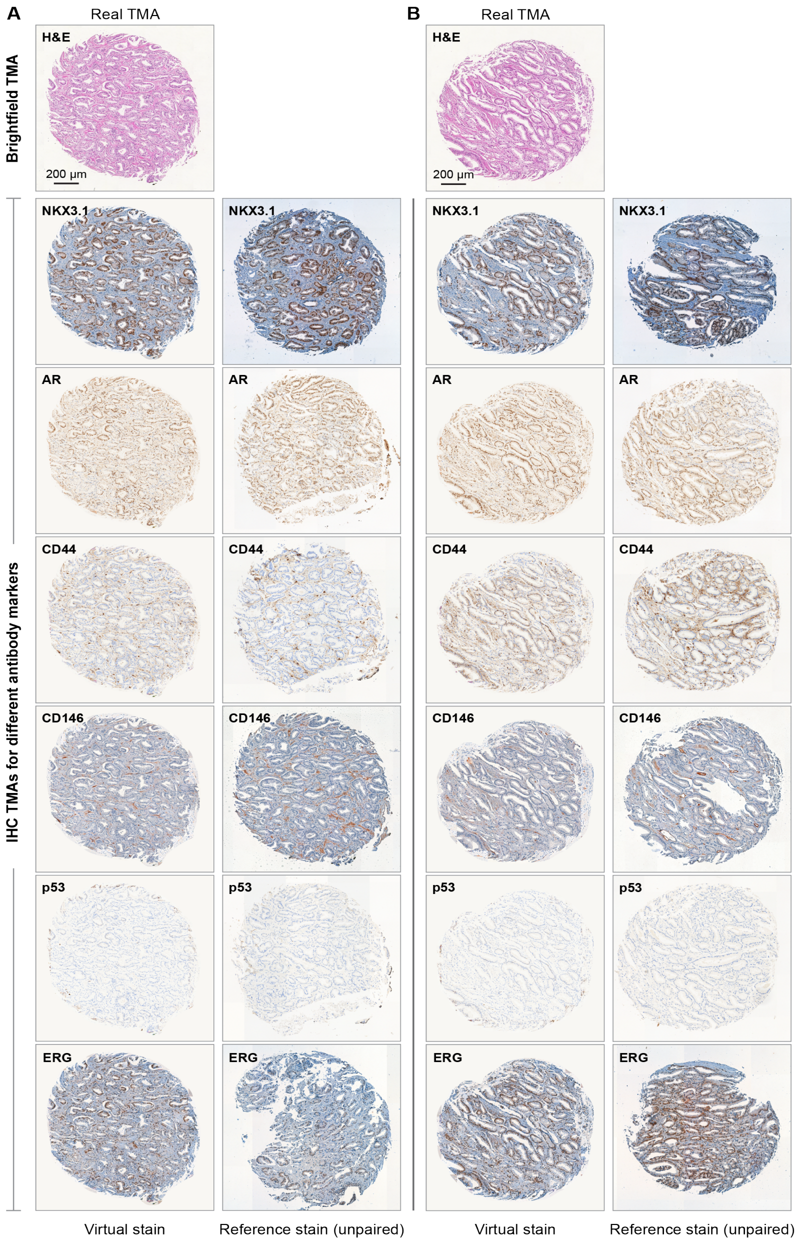


**Fig. A1** Qualitative evaluation of VirtualMultiplexer for two TMAs in the EMPaCT dataset, additional to the qualitative samples presented in Figure 3. **(A)** and **(B)** present two H&E stained TMAs and corresponding virtual stainings for six IHC markers. Columns two and four present reference IHC stained TMAs for the same core.

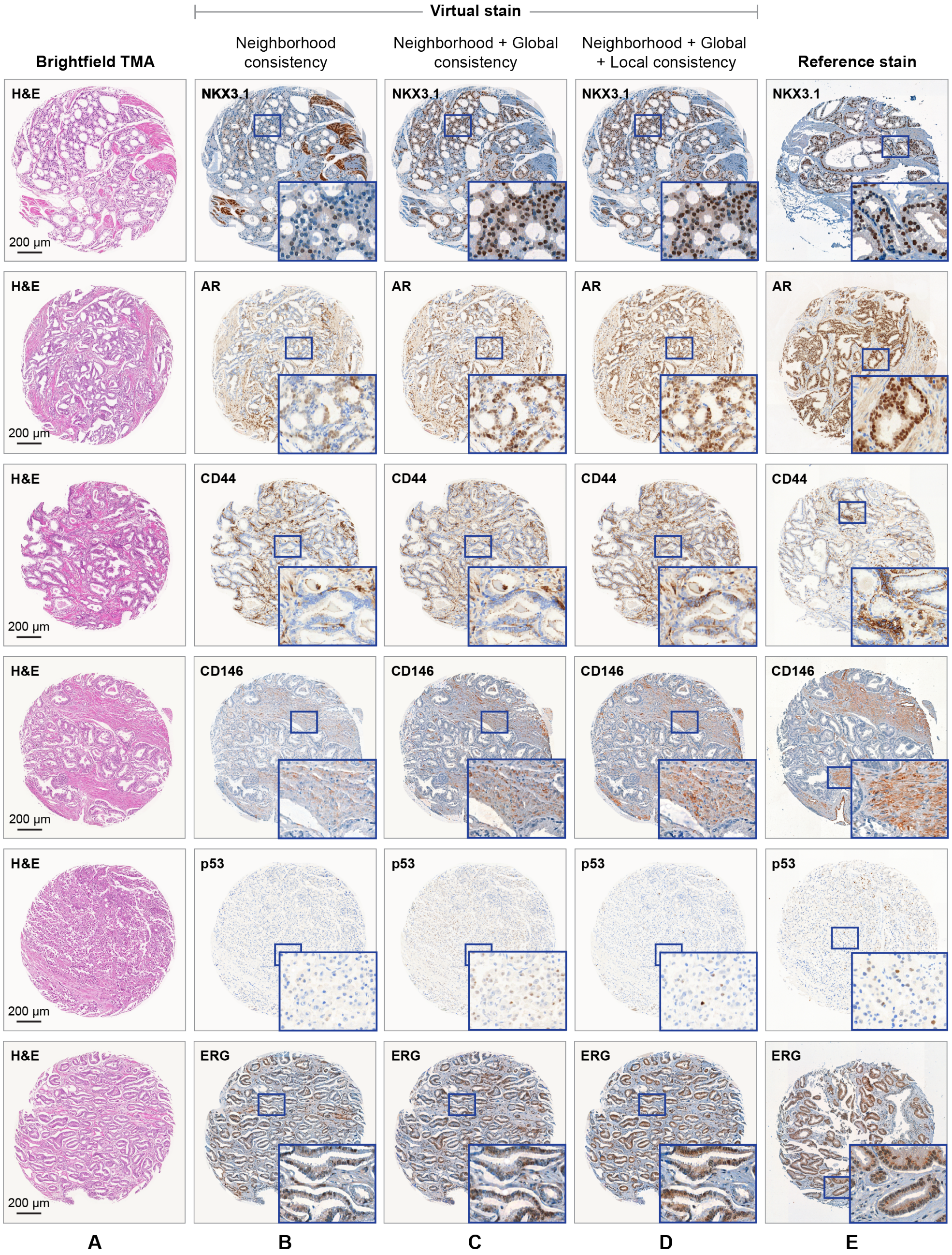


**Fig. A2** Qualitative evaluation of the impact of multi-scale consistency objectives on the virtual staining quality of the VirtualMultiplexer across six IHC markers, presented in each row. **(A)** Sample H&E cores from the EMPaCT dataset. **(B)**, **(C)**, **(D)** Corresponding virtually stained IHC cores for training the VirtualMultiplexer with neighborhood consistency, neighborhood and global consistencies, and neighborhood, global, and local consistencies, respectively. The bounding boxes highlight zoomed-in regions in the IHC cores. **(E)** Reference IHC cores corresponding to the cores in **(A)**.

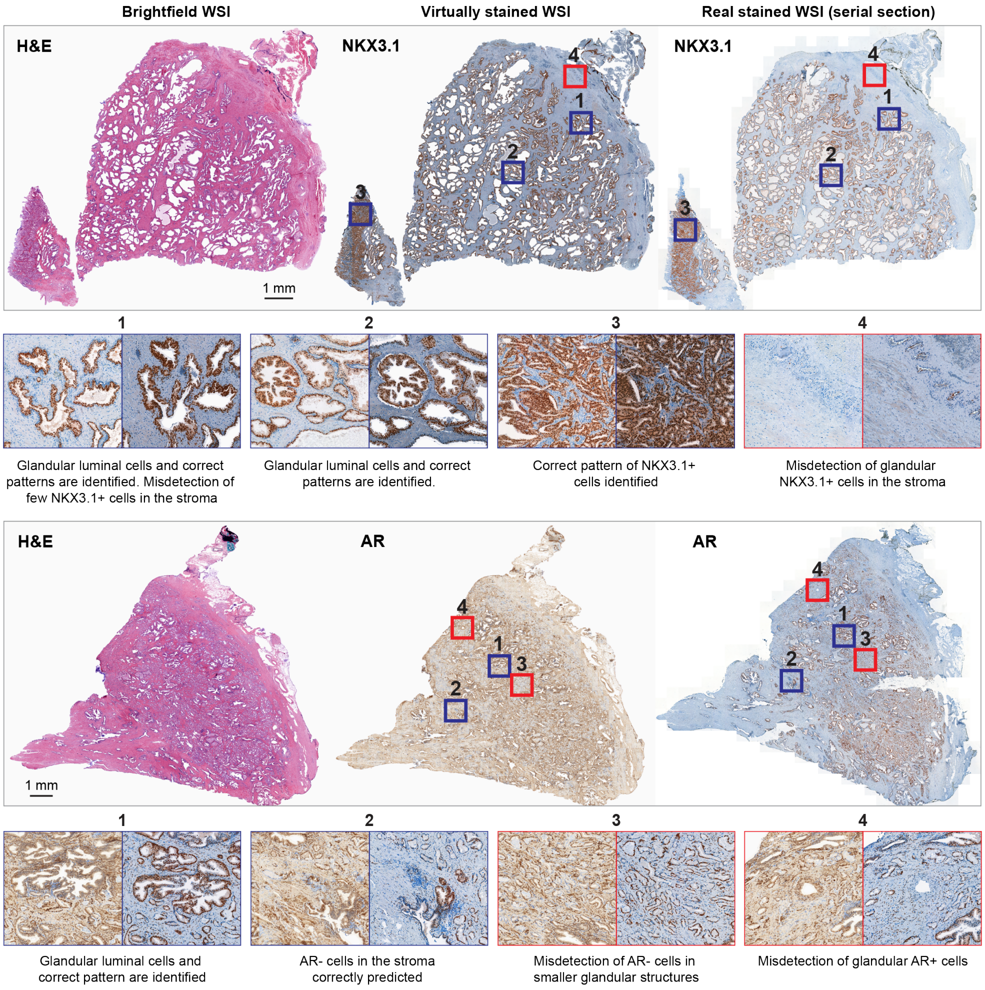


**Fig. A3** Transfer learning from TMAs to WSIs of prostate cancer tissue, additional to the qualitative samples presented in Figure 5. Example of H&E (left image), virtual IHC (middle image), and real IHC (right image) staining for NKX3.1 (top) and AR (bottom) of prostate cancer tissue WSIs. Blue-framed zoomed-in regions display accurate staining pattern. Red-framed zoomed-in regions display examples of virtual staining mispredictions.

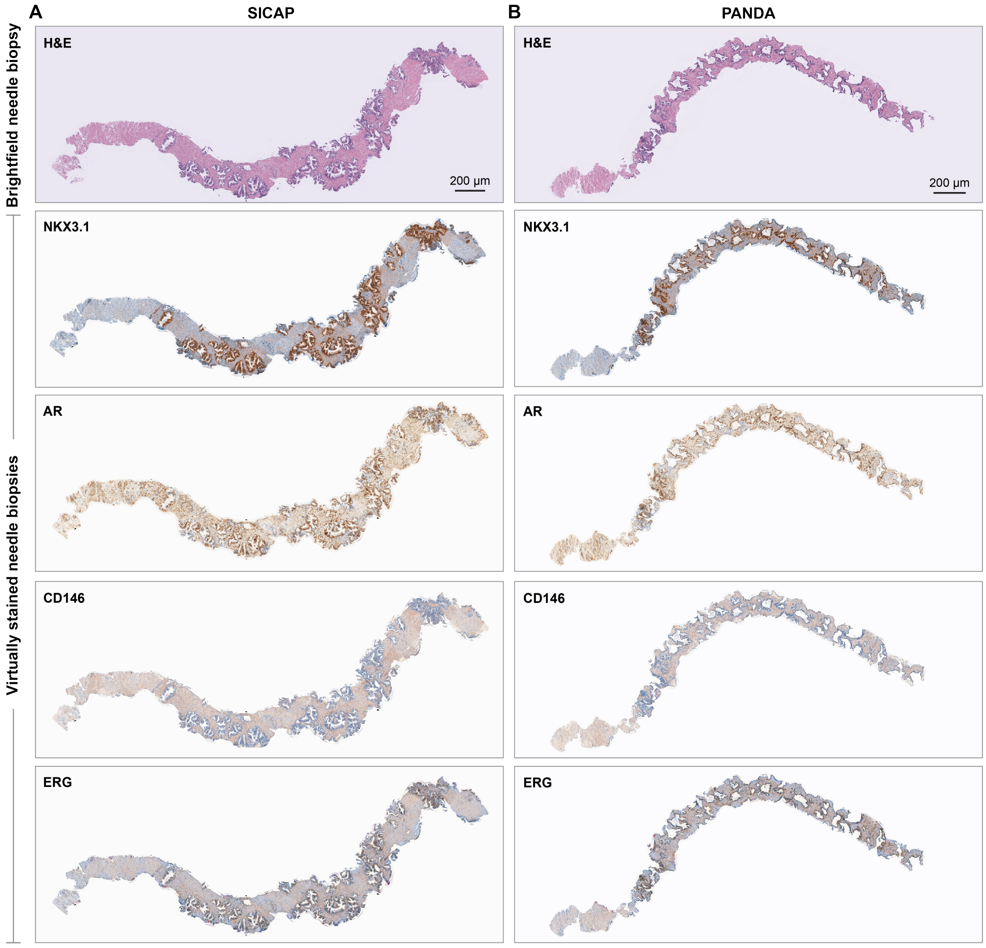


**Fig. A4** Transfer learning from TMAs to needle biopsies of prostate cancer tissue, additional to the qualitative samples presented in Figure 7. **(A)** and **(B)** present H&E biopsies from SICAP and PANDA datasets, respectively, and corresponding virtually stained IHC biopsies for six markers.

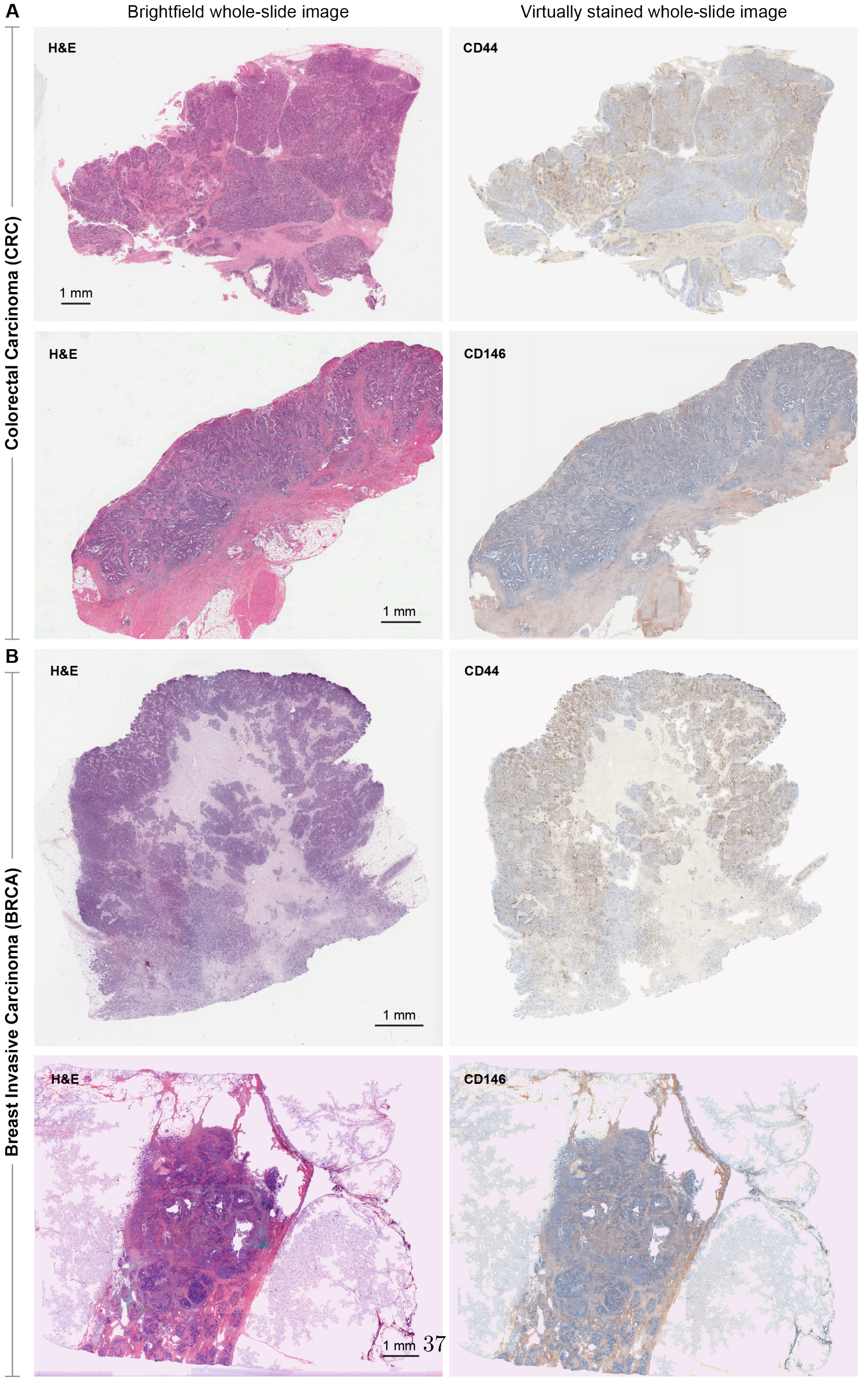




**Fig. A5** Transfer learning from TMAs to WSIs of different tissue types from TCGA cohort. **(A)** H&E WSIs and **(B)** corresponding virtually stained IHC WSIs from colorectal carcinoma (top two rows) and breast invasive carcinoma (bottom two rows). For both the tissue types, the virtual stainings are produced for relevant CD44 and CD146 IHC markers.

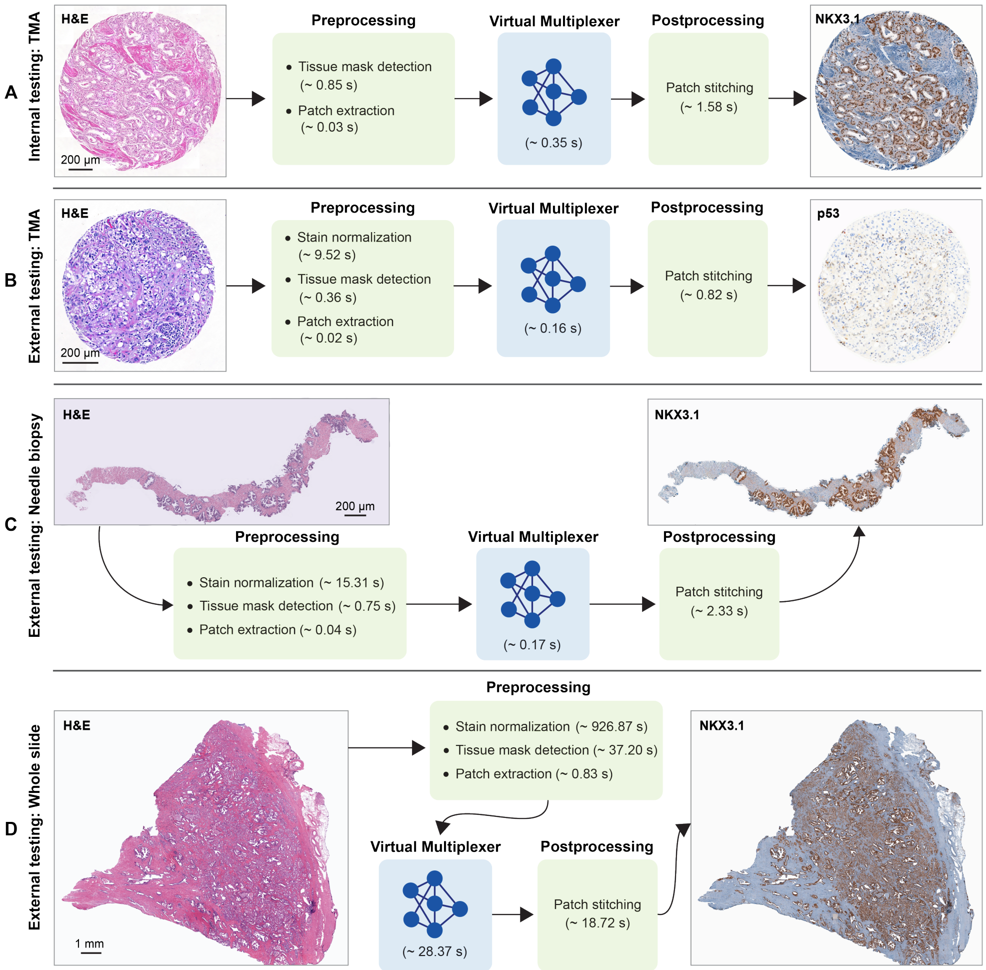


**Fig. A6** Runtime estimation of all components, *i.e.*, , preprocessing, virtual staining, and post-processing, in the proposed computational workflow. **(A)**, **(B)**, **(C)** and **(D)** present the runtime estimations for an in-domain TMA from the EMPaCT dataset, an out-of-domain TMA from the PDAC dataset, an out-of-domain needle biopsy from the SICAP dataset, and an out-of-domain WSI from the in-house dataset. Noticeably, the most time-consuming component in the workflow is the stain normalization, which is performed for all out-of-domain scenarios. This step mitigates the H&E staining variability between the training EMPaCT cores and the test out-of-domain sample.
